# Tracking the evolutionary trajectory of a young hybrid plant pathogen

**DOI:** 10.1101/2025.10.10.681615

**Authors:** Jigisha Jigisha, Urszula Piechota, Paweł Czembor, Szilvia Bencze, Mónika Cséplő, Marta S. Lopes, Thomas Miedaner, Philipp Schulz, Fabrizio Menardo

## Abstract

A common mechanism by which emerging plant pathogens gain the ability to infect new hosts is hybridization. Despite its widespread occurrence, the outcomes of hybridization remain largely unpredictable within current evolutionary frameworks, making empirical studies key for identifying patterns and establishing principles underlying hybrid evolution. Here, we track the evolutionary trajectory of *Blumeria graminis forma specialis triticale* (*B.g. triticale*), or triticale powdery mildew, a pathogen that emerged recently through hybridization of two powdery mildew forms specialized on wheat (*B.g. tritici*) and rye (*B.g. secalis*). Using a genomic dataset of 652 isolates from the three *formae speciales*, we show that they persist as three isolated lineages. Results from our infection assays and sampling suggest that *B.g. triticale* is not well adapted to infect wheat under field conditions, indicating that isolation between *B.g. tritici* and *B.g. triticale* may be maintained by partial niche separation. We investigated the genomic changes following hybridization and found that at least a third of the hybrid genome is fixed for *B.g. tritici* ancestry in contemporary populations, and around 1% is fixed for the *B.g. secalis* ancestry. We identified several loci to be under recent positive selection in *B.g. triticale*, most of which coincide with known genes and regions of fixed or nearly fixed local ancestry. Overall, we highlight the role of ecological isolation in preventing gene flow between *B.g. triticale* and its parental lineages. We reveal the rapid stabilization of its genome after hybridization and show that some of these changes were likely shaped by selection.

**Significance statement:** Emerging plant diseases pose serious risks to biodiversity and food production. A common mechanism by which fungal plant pathogens emerge on previously uncolonized hosts is hybridization. Hybridization brings together novel genetic combinations from different lineages that can facilitate adaptation to new environments. However, the evolutionary fate of hybrid pathogens is difficult to predict, and empirical studies of natural populations can help shed light on the dynamics of pathogen evolution following hybridization. Here, we use genomic data and infection assays to follow the evolutionary trajectory of a recently emerged hybrid plant pathogen. We show that the hybrid persists as an independent lineage distinct from both parents and analyze the genomic changes that have occurred in the few decades since initial hybridization.

## Introduction

Rapid developments in industrial agriculture and global trade networks, coupled with climate change, provide fertile ground for the emergence of plant diseases responsible for significant losses in biodiversity and food production (Bebber 2015; Savary et al. 2019; Corredor-Moreno & Saunders 2020; Ristaino et al. 2021). Emerging plant pathogens are characterized by an increase in incidence, changes in pathogenicity, expansion of geographic range, or colonization of new hosts (Anderson et al. 2004). The appearance of diseases on novel hosts is especially concerning, as there is no shared evolutionary history between host and pathogen, and this can lead to outbreaks of greater severity (Woolhouse et al. 2005). Research investigating the genetic underpinnings of such epidemics has highlighted the role of processes such as mutation, horizontal gene transfer, recombination and hybridization in facilitating host shifts and range expansion (reviewed in Corredor-Moreno & Saunders 2020).

Hybridization is particularly relevant in the context of fungal plant pathogens. By bringing together a novel mix of genotypes from distinct lineages, hybridization, both somatic and sexual, can increase the evolutionary potential of a pathogen and facilitate colonization of previously unoccupied niches (Brasier 2000; Arnold 2004; Depotter et al. 2016; Stukenbrock 2016; Li et al. 2019; Peck et al. 2024). The frequency of successful hybridization in nature depends on several factors, including, but not limited to, the extent of habitat overlap between populations (sympatry, allopatry or parapatry), the time since their divergence, and the presence of pre- and post-zygotic barriers in sexually reproducing organisms (Barton 2001; Taylor & Larson 2019; Steensels et al. 2021). Over evolutionary time, the outcomes of hybridization can be manifold. Hybridization may lead to the formation of new, stable pathogen species through (i) allopolyploidy, such as in the case of *Botrytis alii* (onion rot) (Nielsen & Yohalem 2001) and *Verticillium longisporum* (Verticillium stem striping) (Clewes et al. 2008; Inderbitzin et al. 2011), or through (ii) homoploidy if gene flow is restricted between hybrid and parental populations like in *Zymoseptoria pseudotritici* (Stukenbrock et al. 2012). In some cases, the hybrids may outcompete one or more parental populations and replace them completely. This could explain why parental lineages of several known hybrid pathogens, like *Phytophthora andina* (Goss et al. 2011) and *V. longisporum* (Depotter et al. 2016) have not been sampled and remain undetermined. An alternate outcome of hybridization is introgression, as observed between diverged lineages of *Hemileia vastatrix*, the pathogen responsible for coffee leaf rust (Silva et al. 2018), and between different *Ophiostoma* species causing Dutch elm disease (Hessenauer et al. 2020). In such cases, hybrids exist as ‘transient intermediates’ and facilitate, through successive rounds of backcrossing, the transfer of genomic fragments from one parent to the other.

As a result of the complex interactions involved in hybridization, the genomes of hybrids are extremely dynamic (Runemark et al. 2019). Populations that have recently undergone, or those that are currently undergoing, hybridization (e.g. in hybrid zones), can act as “windows into the evolutionary process” (Harrison 1990), and genomic data from these populations can reveal intricate details about adaptation, diversification and the onset of speciation (Nolte & Tautz 2010; Gompert et al. 2017). A powerful approach for analyzing such data is to track variation in local ancestry along the genome, among individuals, and across time (Gravel 2012; Moran et al. 2021). Such methods have been used in various taxonomic groups to yield insights into how the fundamental evolutionary forces of drift, recombination, selection and gene flow affect hybrid populations (Leitwein et al. 2018; Schumer et al. 2018; Chaturvedi et al. 2020; Howard-McCombe et al. 2023; Nevado et al. 2024). A notable example involving fungal plant pathogens was reported in a study on *Pyricularia oryzae*, the causal agent of blast and leaf spot diseases of numerous crops and grasses (Rahnama et al. 2023). The authors showed that host-range expansion of this pathogen to include wheat and ryegrass was facilitated by recent hybridization among *P. oryzae* populations specialized on other grass species. This study highlighted the relative importance of repartitioned standing variation over new mutations in the establishment of pathogen populations on novel hosts.

Another remarkable instance of emergence of a novel plant disease through sexual hybridization comes from *Blumeria graminis* (*sensu stricto)* (Liu et al. 2021), the obligate-biotroph ascomycete responsible for the powdery mildew disease in various cereals and grasses. This species is further classified into different *formae speciales* (*ff. spp.*), that do not differ from each other morphologically but are specialized to infect distinct host species. As a result of rarely encountering individuals occurring on non-hosts in nature, gene flow among the *ff. spp.* is restricted, and they form distinct genetic groups (Menardo et al. 2017). Two of these diverged forms, *B.g. tritici* and *B.g. secalis*, adapted to infect wheat and rye respectively, were shown to have recently undergone hybridization (Menardo et al. 2016). This led to the formation of a new homoploid *forma specialis*, *B.g. triticale*, which is capable of infecting the previously immune cereal triticale. Triticale, itself the result of artificial hybridization between wheat and rye, was first used in agriculture in the 1960s and was known to be resistant to powdery mildew until 2001, when the first infections were observed in the field (Mascher et al. 2006; Walker et al. 2011). Powdery mildew has since then become a major disease of triticale in Europe. Genomic analyses revealed *B.g. triticale* to be a mosaic of alternating genomic segments inherited from *B.g. tritici* and *B.g. secalis* (Menardo et al. 2016; Müller et al. 2021). Infection tests in the laboratory have indicated that in addition to infecting triticale, *B.g. triticale* can also grow on wheat, suggesting that there may be overlaps in the ecological niches of the hybrid and at least one of the parental lineages (*B.g. tritici*).

Here, we utilized this unique system of a very recently emerged (50 to 25 years ago) plant pathogen, *B.g. triticale* (Menardo et al. 2016), to investigate its evolutionary trajectory following hybridization. Using a dataset of 652 genomes from three *ff. spp.* (tritici, secalis and triticale), sampled at different time points before and after disease emergence on triticale, we aimed to (i) characterize the degree of isolation between the hybrid and parental populations, (ii) identify potential factors affecting gene flow between the *ff. spp.*, and (iii) assess the extent of variation in local ancestry along *B.g. triticale* genomes over space and time to study the genomic changes that accompany the establishment of a stable hybrid lineage. We found that *B.g. triticale* forms a lineage isolated from both parents, and ecological separation likely contributes to limit gene flow between the three *ff*. *spp.* We also found that at least a third of the hybrid genome was fixed for the *B.g. tritici* ancestry in contemporary populations of triticale powdery mildew, and that the few regions that were fixed for the *B.g. secalis* ancestry constituted around 1% of the genome and were restricted to specific chromosomes. Further, we identified several loci that were putatively under recent positive selection, most of which overlapped with regions with fixed or nearly fixed local ancestry, and with previously annotated genes, including two candidate effectors. Finally, we found that the mosaic pattern of local ancestry along the *B.g. triticale* genome is largely stable between isolates collected in different periods and locations, and that on average *B.g. secalis*-origin and *B.g. tritici*-origin segments make up roughly 17% and 83% of the hybrid genome respectively.

Overall, our results show that a large proportion of the *B.g. triticale* genome stabilized rapidly, within one to a few decades after hybridization, and highlight the role of ecological isolation in limiting gene flow between this hybrid pathogen and its parental lineages.

## Results

### Sampling and determination of *formae speciales*

A total of 326 *Blumeria graminis* isolates were obtained from over 90 locations in Europe and surrounding regions in 2022 and 2023 (see Methods for details; Table S1) (Jigisha et al. 2025). Since the *formae speciales* cannot be distinguished morphologically, and because there may be overlaps in their niches in nature, we performed host specificity assays and genomic analyses to confirm the identity of these isolates. The panel of hosts tested included the susceptible wheat cultivar Kanzler, two triticale cultivars, Bedretto and 1011.58329 (Agroscope), and a rye cultivar, Matador (Fig. 1a, Table S1). We found that 276 isolates belonged to *B.g. tritici*; these have been described previously and were included in the dataset analyzed in Jigisha et al. 2025. All *B.g. tritici* isolates could successfully infect Kanzler, 150 could grow on Bedretto to varying degrees, and none could infect the other triticale cultivar (Agroscope), or Matador. Two isolates were able to infect rye, but not wheat or triticale, and were classified as *B.g. secalis*. The remaining 48 isolates were able to grow on wheat and both triticale cultivars, but not on rye. We classified these isolates as triticale powdery mildew, or *B.g. triticale*.

**Fig. 1.**
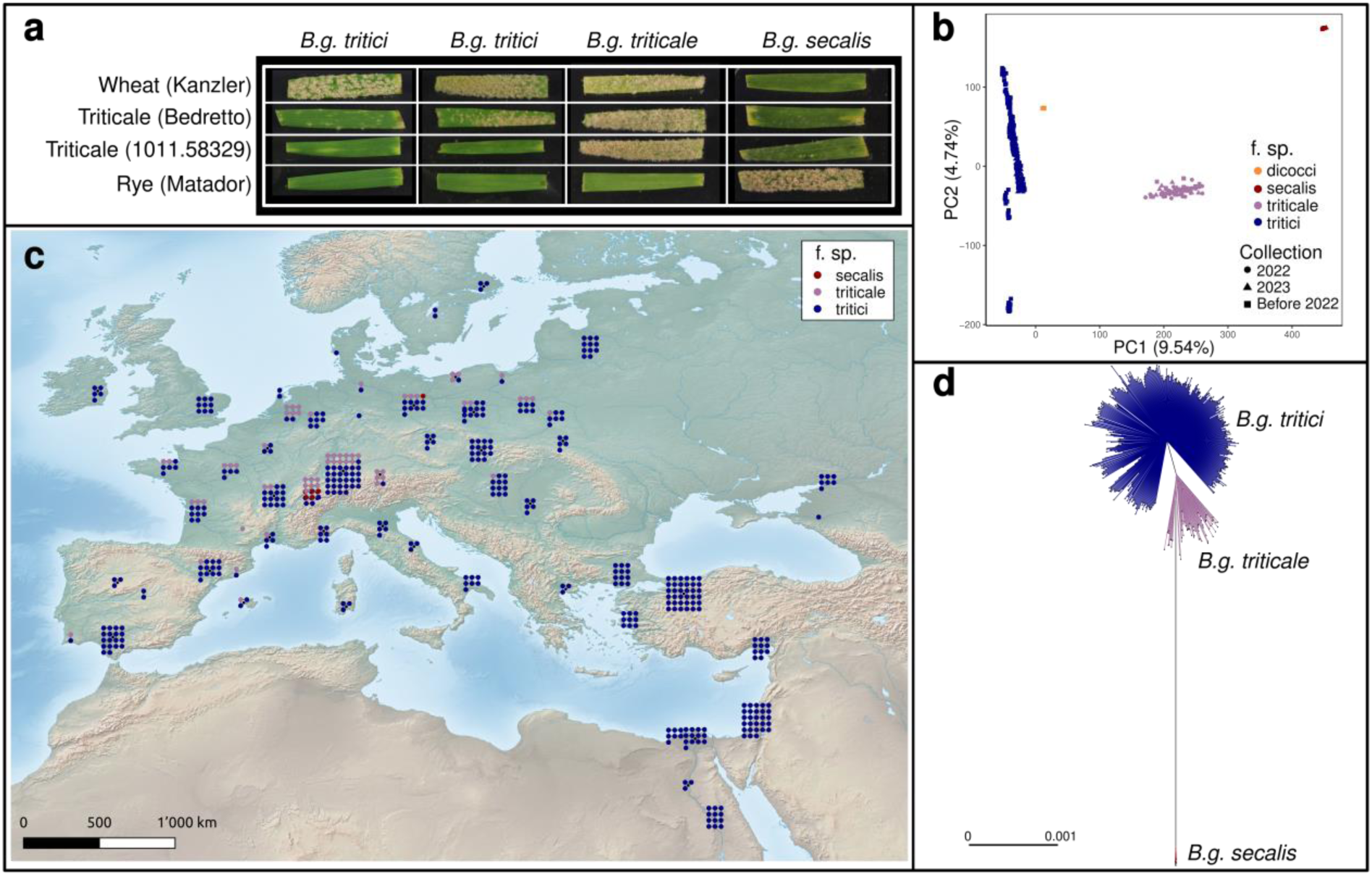
*Formae speciales* differ in host specificity and form distinct genetic groups. **(a)** Representative images from the host-specificity assays. Rows correspond to the host cultivar tested. Each column shows the infection results for one isolate. The isolates shown here are: (L-R) ESCT052207, ITBO062201, PLRA092317 & DEBD062355. Only one of the three biological replicates (per host and per isolate) is shown here. **(b)** PC1 vs PC2 of the PCA performed on 2,431,821 SNPs from 652 isolates sampled globally. Each point represents one isolate and the colours represent the *forma specialis*. The colour scheme is the same in panels **b-d**. **(c)** Map showing sampling locations of the 484 isolates of different *ff. spp.* collected from Europe and neighbouring regions between 1980 and 2023. (d) Phylogenetic network of 484 isolates of different *ff. spp.* based on pairwise genetic distances.

We confirmed this classification with genomic data. We combined the genome sequences of all the isolates described above with publicly available genomes of *B.g. tritici*, *B.g. secalis*, *B.g. dicocci* (a *f. sp.* occurring in the Middle East and preferentially infecting tetraploid wheat; Menardo et al. 2016; Menardo et al. 2017; Ben-David et al. 2016; Sotiropoulos et al. 2022), and *B.g. triticale*.

Following filtering, the final dataset contained 652 isolates (see Methods for details). All genome sequences were mapped to the *B.g. tritici* reference genome 96224 (see Methods). It has been shown previously that the genomes of these *formae speciales* are highly collinear and show > 98% homology with the 96224 reference assembly (Müller et al. 2019; Sotiropoulos et al. 2025). In our dataset, the median proportion of reads mapped to the *B.g. tritici* reference genome was greater than 98% for all *ff. spp*. Upon variant calling, we obtained 2,431,821 biallelic SNPs across 652 isolates that were used to perform a principal component analysis (PCA). The first two principal components explained 14.28% of the total variation and grouped samples into four distinct clusters corresponding to the four *formae speciales* – dicocci, tritici, secalis and triticale (Fig 1b). The two new *B.g. secalis* isolates clustered with the old *B.g. secalis*, and the 48 isolates that were phenotypically characterized as *B.g. triticale* clustered together with the older collection of *B.g. triticale* samples. Of the total 70 *B.g. triticale* genomes identified from this PCA, we retained 62 non-clonal isolates. Furthermore, we excluded all dicocci isolates, and tritici isolates collected outside of Europe and the Middle East, resulting in a dataset of 484 isolates for downstream analyses (see Methods; Table S1).

### *B.g. triticale* is an isolated lineage

From the results of the host-specificity assays and the PCA described above (Fig. 1a-b), it is evident that *B.g. triticale* forms a group with phenotypic and genetic features distinct from its parental *formae speciales*. Taken together with the fact that we sampled both *B.g. tritici* and *B.g. secalis* from the field as recently as 2023, this suggests that the hybrid has neither merged with nor replaced either of the parental forms yet. Moreover, isolates sampled in 2022-2023 and in previous periods grouped together in the PCA according to their *f. sp.*, indicating that the genetic structure of these three *ff. spp.* has remained stable in the last ten years.

To further explore the extent of isolation among the three *formae speciales* in Europe, we first constructed a phylogenetic network of 484 *Blumeria graminis* isolates sampled in Europe and surrounding regions (Fig. 1c). This network, based on pairwise genetic distances, clearly differentiated *B.g. tritici*, *B.g. secalis* and *B.g. triticale* from each other (Fig. 1d). Moreover, as also previously reported in Menardo et al. 2016 and seen from the PCA described above, *B.g. triticale* was genetically intermediate between *B.g. tritici* and *B.g secalis*. Next, we characterized the degree of shared ancestry between these 484 isolates using fineSTRUCTURE (Lawson et al. 2012), a method which accounts for linkage between SNPs and uses haplotype similarity to identify population structure. All individuals belonging to the same *forma specialis* shared more ancestry with each other than they did with individuals from the other groups (Fig. S1). This method also broadly recovered the population subdivisions in *B.g. tritici* described previously (5 populations - N_EUR, S_EUR1, S_EUR2, ME, TUR) (Jigisha et al. 2025). Additionally, it revealed the substructure present within *B.g. triticale*, identifying two main groups: one comprising isolates sampled mainly from the eastern parts of Europe, and the other, mostly from the west. Further, this analysis showed that the *B.g. triticale* isolates shared more genetic similarity with *B.g. tritici* isolates sampled from Northern Europe compared to those from Southern Europe and the Middle East. This was also confirmed by the population trees constructed using TreeMix (Pickrell & Pritchard 2012) (Fig. S2) and is consistent with the hypothesis that *B.g. triticale* originated in Northern Europe (Menardo et al. 2016).

It was previously shown that the genome of *B.g. triticale* consists of a mosaic of large segments with alternating *B.g. secalis* and *B.g. tritici*-like genotypes and that *B.g. tritici-*like segments made up more than 80% of the *B.g. triticale* genome (Menardo et al. 2016). This was interpreted as the result of a series of backcrosses following the initial hybridization event between wheat and rye powdery mildew. If isolation among *B.g. tritici* and *B.g triticale* was weak and gene flow had since continued, we would expect to find the introgression of genomic segments with a *B.g. secalis* genotype into *B.g. tritici*. To investigate this, we examined the genomes of 255 *B.g. tritici* isolates sampled in 2022-2023 for signs of recent *B.g. secalis* introgression. Using polymorphisms fixed for different alleles (substitution sites; see Methods) between *B.g. secalis* and the older *B.g. tritici* population comprising individuals sampled until 2001, we assigned parental origin to genomic segments in each of the 255 recent *B.g. tritici* isolates. We did not find evidence for large introgressions of *B.g. secalis* genotype segments in *B.g. tritici*, and on average, the *B.g. secalis* origin segments covered 0.076% of the genomes, with some variation among isolates belonging to different populations (Fig. S3, Appendix S1). Furthermore, comparing the genomic composition of the *B.g. triticale* individuals sampled in the past (2009-2013) with those sampled recently (2022-2023), we found that the proportion of genome covered by *B.g. secalis* origin segments did not differ between the two groups and was about 17% on average (Welch’s t-test p = 0.74; Fig. S4).

Finally, we constructed phylogenetic trees for three of the genes associated with the two mating types in *Blumeria graminis* (Wicker et al. 2013). This analysis showed no signs of new *B.g. secalis* introgression into *B.g. triticale* in the last ten years (Appendix S2, Fig. S5). Combining the results from all the analyses described above, we conclude that *B.g. tritici*, *B.g. secalis* and *B.g. triticale* persist as independent isolated lineages, with no evidence of substantial gene flow between them.

### Factors maintaining isolation between *B.g. tritici* and *B.g. triticale*

Laboratory crosses between *B.g. tritici* and *B.g. triticale* have been shown to result in viable offspring (Müller et al. 2019). It is therefore intriguing that *B.g. triticale* still forms a lineage isolated from *B.g. tritici*. In this section, we attempt to identify factors that may be maintaining isolation between these two *formae speciales* in nature.

The infection tests described above and previously (Menardo et al. 2016) have shown that *B.g. triticale* and *B.g. tritici* can both grow on wheat and therefore might have the opportunity to meet and reproduce in nature. However, these tests were performed on susceptible wheat varieties and under ideal conditions for pathogen growth. Whether the two *ff. spp.* effectively share the same niche in nature and infect the same hosts remains unknown. Our sampling data provided a first indication. In 2022 and 2023 we sampled *B. graminis* isolates with two approaches: from infected wheat fields and from trap plants of susceptible wheat varieties (Methods). In both cases we did not know *a priori* whether the samples belonged to *B.g. tritici* or *B.g. triticale*. Considering only samples originating from the core geographic range of *B.g. triticale*, (Northern Europe including Northern Spain), only three of the 118 samples obtained from wheat fields belonged to *B.g. triticale* (2.5%); conversely, 11 of the 87 samples obtained from trap wheat plants were classified as *B.g. triticale* (12.6%). The significant underrepresentation of *B.g. triticale* samples from wheat fields (Fisher exact test *p* < 0.01) suggests that *B.g. triticale* does not grow particularly well on wheat in field conditions. To further investigate differences in the ecological niches of *B.g. tritici* and *B.g. secalis* we performed infection assays on 20 wheat lines carrying a single, or a combination of, known powdery mildew resistance (*Pm*) genes (Methods; S2 Table). A principal component analysis based on the results for all the tested isolates (34 *B.g. triticale* and 104 *B.g. tritici*) highlighted that the virulence profiles of the two *ff. spp.* were distinct (Fig. 2). We found that on average *B.g. triticale* isolates were equally or less virulent than *B.g. tritici* isolates on the 20 tested wheat lines, with the only exception being the line carrying *Pm35* (Fig. S6). Moreover, some of the largest differences were observed for *Pm* genes that have been used prominently in wheat breeding, such as *Pm2*, *Pm4a*, and *Pm8* (Sánchez-Martín & Keller 2019), a pattern consistent with a previous report (Troch et al. 2013).

**Fig. 2.**
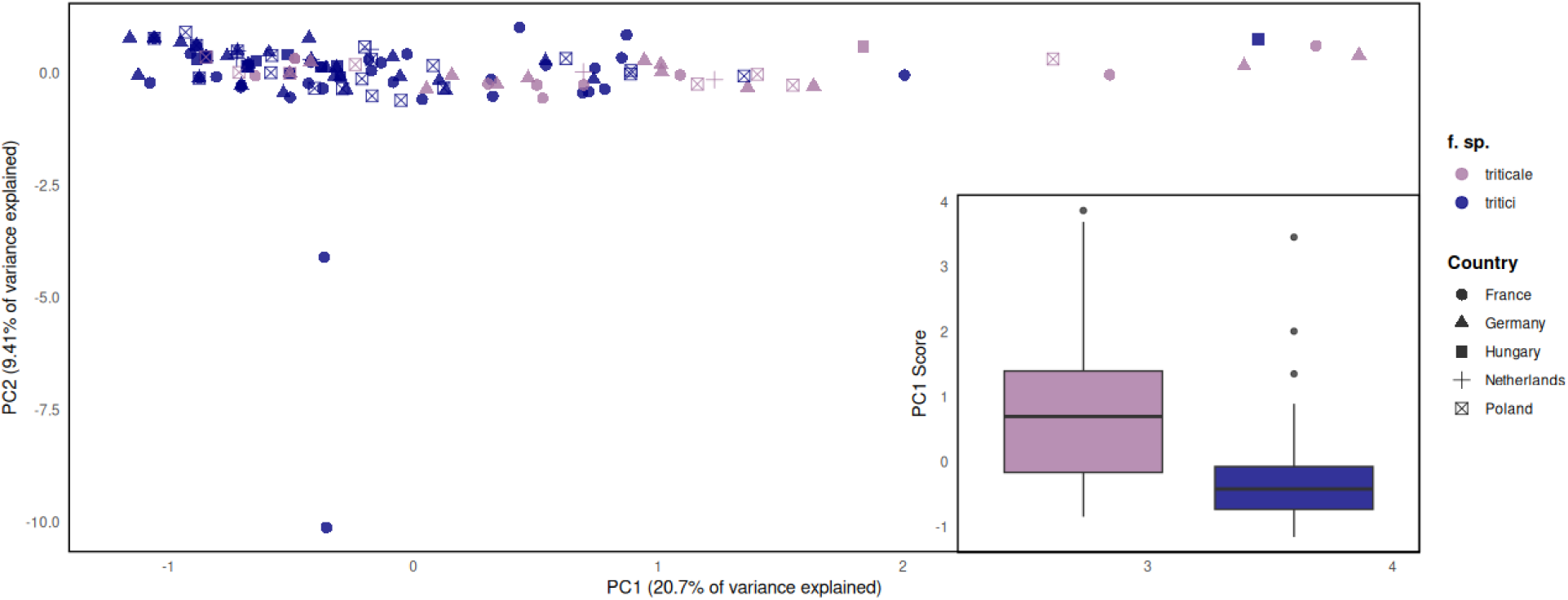
*B.g. tritici* and *B.g. triticale* have distinct virulence profiles. Principal component analysis based on the infection tests performed for *B.g. tritici* and *B.g. triticale* isolates on 20 wheat lines carrying different *Pm* genes. Each point represents one isolate. The boxplot in the inset compares the distribution of the PC1 scores for isolates belonging to different *ff. spp*. The horizontal bars depict the median.

Our sampling results, together with the outcome of the infection assays, suggest that these two *formae speciales* have only limited opportunity to encounter each other in nature, as *B.g. triticale* is not well adapted to infect wheat under field conditions. These findings support those of Czajowski and Czembor (Czajowski & Czembor 2016), who reported that *B. graminis* isolates sampled from triticale were, on average, less virulent on wheat than those sampled on wheat. Nevertheless, while there are differences in the virulence profiles of *B.g. tritici* and *B.g. triticale*, there are also overlaps, and there might be other mechanisms, in addition to ecological separation, involved in the maintenance of isolation between these two lineages.

Mating of two diverged lineages may bring together a combination of alleles that experience negative epistatic interactions; alleles that are benign or beneficial in one genomic background can have deleterious effects in alternate backgrounds. The accumulation of such between-locus incompatibilities, known as Dobzhansky Muller incompatibilities (DMIs) (Dobzhansky 1937; Muller 1942), can lead to postzygotic isolation between populations. We identified possible DMIs in hybrids between *B.g. tritici* and *B.g. triticale* using the genome sequences of 116 progenies of a laboratory cross between 96224 (*B.g. tritici*) and THUN-12 (*B.g. triticale*) described in Müller et al. 2019 (see Methods). We found 333 pairs of genomic windows where both regions, present on different chromosomes, were inherited from the same parent significantly more often than from different parents (Fig. S7, Table S3). In other words, these regions did not segregate independently in the offspring, making them candidates for DMIs. Such regions were found on all chromosomes except chromosome 11 and covered ∼ 7.84% of the whole genome. In 27 (of 333; 8.1%) identified putative DMI pairs, one genomic region of the pair showed *B.g. secalis*-like genotype in THUN-12, and there was no pair where both regions could be assigned to *B.g. secalis* origin (Fig. S7), suggesting that these putative DMIs are unlikely to be caused by diverged genotypes of *B.g. tritici* and *B.g. secalis*. Nonetheless, 16 of the 186 unique regions implicated in putative DMIs overlapped with genomic windows associated with high divergence between the *B.g. triticale* and *B.g. tritici* N_EUR populations (d_xy_, F_ST_ outliers), and 14 of these corresponded to *B.g. secalis* segments in the genome of the parental isolate THUN-12.

Altogether, our findings suggest that *B.g. triticale* does not infect wheat very often in field conditions, and therefore *B.g. tritici* and *B.g. triticale* experience some degree of ecological separation in nature, which can contribute towards maintaining isolation between them. However, ecological isolation is not complete, and there might be additional factors limiting gene flow between these two *ff. spp*. One of such factors could be the presence of DMIs, but whether these contribute to the isolation of wheat and triticale powdery mildew remains to be tested.

### At least a third of the *B.g. triticale* genome is fixed for *B.g. tritici* ancestry

In the immediate generations following hybridization, hybrid populations show high amounts of variation in local ancestry among genomes of different individuals. If hybridization is ongoing, this variation in ancestry proportions is expected to remain high (Runemark et al. 2019). In contrast, as the hybrid gets established as an isolated lineage, recombination breaks down the large ancestry tracts into smaller blocks, local ancestry variation among individuals is expected to decrease and eventually, specific ancestry tracts reach fixation in the hybrid population (Buerkle & Rieseberg 2008). The time taken to genome stabilization can vary from species to species, and the process may be driven by selection, for example against certain ancestry tracts involved in incompatibilities, or favouring those involved in adaptation, or by genetic drift.

To assess genome stability in *B.g. triticale*, we characterized variation in local ancestry for all *B.g. triticale* isolates in non-overlapping windows of 10 Kb along the 11 chromosomes using the substitution-sites method described previously, as well as by employing a nested HMM-based approach, as implemented in MOSAIC (Salter-Townshend & Myers 2019) (Methods, Appendix S3, Table S4). Across the different methods, we covered 72.2%-77.8% of the genome with valid windows (windows with at least one SNP; see Methods; Table S5).

First, we identified the genomic regions that have reached fixation for an ancestry by making use of the 46 *B.g. triticale* isolates sampled in 2022 and 2023. Of the total genome length covered by valid windows, between 33.63% to 46.52% was found to be fixed for one ancestry or the other, depending on the method used (Table S5). The regions fixed for the *B.g. secalis* ancestry were restricted to a few chromosomes across all methods and made up between 0.78% - 1.36% of the total genome covered by valid windows while 32.64%-45.74% was made up of regions fixed for the *B.g. tritici* ancestry, distributed across all 11 chromosomes (Fig. 3, Figs. S8-S9). We also found that some of these regions were fixed already in the earliest collection of triticale mildew isolates (collected in 2009/2010) and have persisted in the population since, while others seem to have risen in frequency to fixation within the last ten years (Fig. S10). Several other genomic windows, though not fixed for an ancestry, showed high average *B.g. secalis* or *B.g. tritici* ancestry across individuals suggesting that these may be on their path to stabilization (Fig. 3, Figs. S8-S9).

**Fig. 3.**
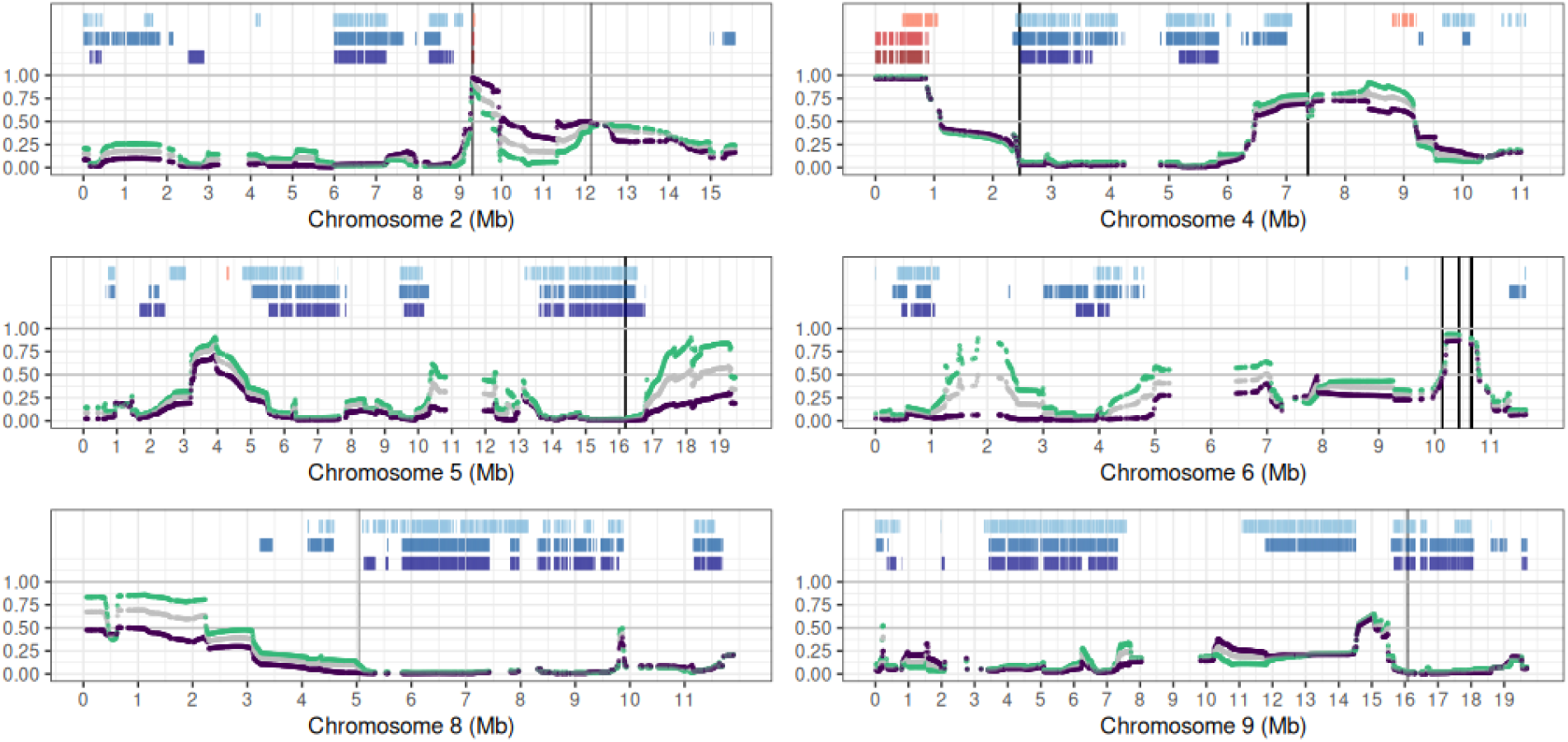
More than a third of the *B.g. triticale* genome is fixed for an ancestry. The blue and red segments at the top of each panel (chromosome) represent the genomic windows fixed for *B.g. tritici* or *B.g. secalis* ancestry respectively across all *B.g. triticale* isolates sampled in 2022-2023. The first row from the top shows results from the substitution sites method, the second row corresponds to the MOSAIC run with old *B.g. tritici* donors and the third belongs to the MOSAIC run with new *B.g. tritici* donors (see Methods and Appendix S3 for details). Plotted in the lower section of each panel is mean *B.g. secalis* ancestry in windows of 10 Kb, as obtained from the MOSAIC run with new donors. Mean ancestry estimates from the alternate MOSAIC run with old donors are plotted in Fig. S9. Mean ancestry across all *B.g. triticale* isolates sampled in 2022-2023 is shown in grey, while green and purple correspond to mean ancestry for the eastern and western populations respectively. The vertical black lines represent loci found to be under recent positive selection from the isoRelate analysis, after FDR correction. Only those chromosomes that house loci under selection are plotted here. For the same figure with all 11 chromosomes, see Fig. S8.

To test if some of these patterns of local ancestry could be explained by selection, we performed genome wide scans for selection based on excess sharing of identical-by-descent segments using all 62 *B.g. triticale* isolates in our dataset. We identified ten regions along the genome that were putatively under recent positive selection (Fig. S11). Four of these regions overlapped with genomic windows that were fixed for an ancestry (Fig. 3), such as those on chromosomes 5 and 9 fixed for *B.g. tritici* ancestry. Interestingly, a region on chromosome 2 that appears to have been fixed for the *B.g. secalis* ancestry only in the last decade was also found to coincide with a peak in the selection scan. Most of the other regions suspected to be under selection were associated with high mean *B.g. secalis* (example, chromosome 6) or mean *B.g. tritici* ancestry (example, chromosome 8).

Overall, our results indicate that a large proportion of the *B.g. triticale* genome has nearly or completely stabilized, and that this was partly due to selection.

### Loci under selection are enriched for functional elements

Fungicides and host resistance genes are common sources of selection acting on agricultural pathogens. In a previous study, genome-wide scans for signatures of selection in *B.g. tritici* populations from Europe and the Middle East revealed several avirulence genes and fungicide targets to be putatively under recent positive selection (Jigisha et al. 2025). We checked if the footprints of selection previously observed in *B.g. tritici* populations were similar to those we detected in *B.g. triticale* in the present study. We found nearly no overlap between the selection scans of *B.g. triticale* and any of the *B.g. tritici* populations (Fig. S12), suggesting that the *formae speciales* experience distinct selection pressures. Further, the loci under selection in *B.g. triticale* did not coincide with any characterized avirulence gene or known fungicide target (Fig. S12, Table S6). However, 7 of the 10 loci identified in the selection scans of *B.g. triticale* were found to overlap with 13 annotated genes in the 96224 *B.g. tritici* reference genome (Table S7, Muller et al. 2019). Two of these 13 genes (BgtE-5850 on chromosome 2 and BgtE-10116 on chromosome 6) have also previously been proposed to encode candidate effectors (small, secreted proteins that facilitate infection by suppressing plant immune responses or modulating host physiology; Lo Presti et al. 2015).

We probed further into the 60 Kb region on chromosome 2 (9290000-9350000) that overlapped with a signature of selection and was fixed for *B.g. secalis* ancestry in the recently sampled isolates of *B.g. triticale* (Fig. 4a). Genome alignment between the 96224 (*B.g. tritici*) and THUN-12 (*B.g. triticale*) reference genomes (Muller et al. 2021) revealed no evidence of major rearrangements in this window (Fig. 4b). In the 96224 genome, this region contained 6 annotated genes, including the candidate effector BgtE-5850. Both BgtE-5850 and the corresponding candidate effector in THUN-12 (BgTH12-03926) are members of E004, the third largest effector gene family in *B.g. tritici*. E004 is a ‘group 1’ effector family, meaning that it encodes short (100-200 aa) and highly expressed proteins (Muller et al 2021), and the previously characterized avirulence gene *AvrPm1a* also belongs to this family (Hewitt et al. 2020).

**Fig. 4.**
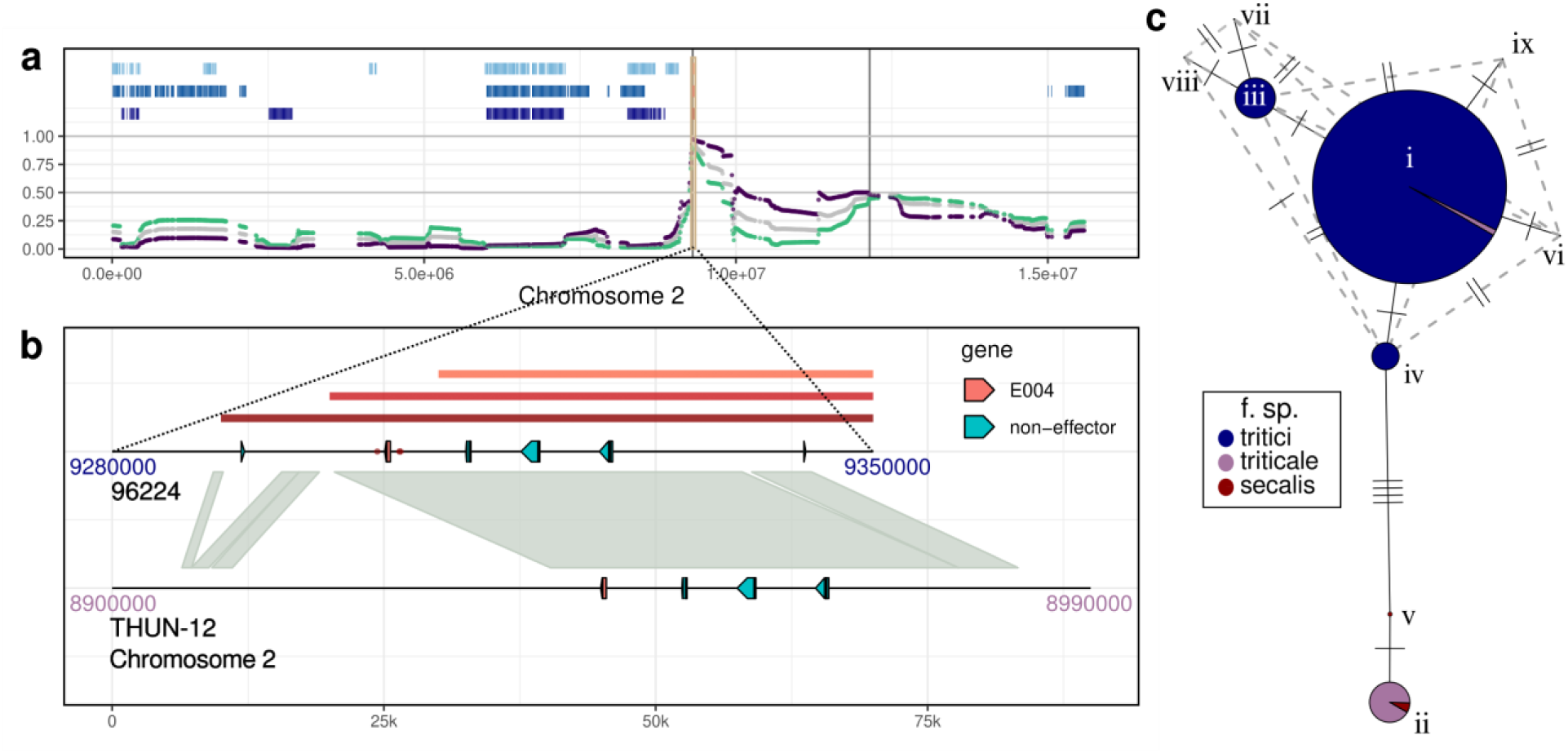
Locus fixed for *B.g. secalis* ancestry contains candidate effector. (**a**) Mean ancestry estimates and regions fixed for an ancestry in *B.g. triticale* on chromosome 2 (same as Fig. 3). (**b**) Zoom-in of the yellow shaded region in **a** on the 96224 genome (coordinates: 9280000-9350000). The red bars on top represent the windows found to be fixed for *B.g. secalis* ancestry in recent *B.g. triticale* isolates using the three different local ancestry estimation methods (top: substitution sites, middle: MOSAIC with old tritici donors, bottom: MOSAIC with new tritici donors). Arrows represent annotated genes. The red dots on either side of the effector represent the SNPs crossing the FDR corrected significance threshold in the *B.g. triticale* selection scan. Alignment of the 96224 genome with the THUN-12 assembly is represented with the synteny plot. (c) Nucleotide haplotype network of BgtE-5850 for 484 *B. graminis* isolates. Each node represents a unique haplotype, as also indicated by the roman numerals. Sizes of the nodes are proportional to the number of isolates displaying that haplotype. Ticks on edges correspond to number of nucleotide differences and dashed grey lines represent alternate connections.

Next, we constructed a nucleotide haplotype network for the BgtE-5850 gene using 484 isolates of the tritici, secalis and triticale *formae speciales*. We found 9 unique haplotypes across all isolates (Fig. 4c; Table S1). Three *B.g. triticale* isolates sampled between 2009-2013 showed the dominant *B.g. tritici* haplotype (haplotype i). All remaining *B.g. triticale* isolates shared the same haplotype as *B.g. secalis* (haplotype ii). Interestingly, we found an alternate *B.g. secalis* haplotype in our dataset (haplotype v) present in two isolates that was absent in the other two *formae speciales*. Overall, our findings on this locus on chromosome 2 demonstrate an example of adaptive introgression from *B.g. secalis* in *B.g. triticale*.

### Tracking variation in local ancestry over space and time

In addition to the genomic regions that are stable, those that show high variation in local ancestry over space and time can also shed light on the processes shaping hybrid genome evolution. We first investigated variation in mean local ancestry among isolates originating from different sampling locations. Population structure analyses of *B.g. triticale* in Europe indicated the presence of two distinct genetic groups with different geographic ranges: one population comprising isolates mainly from eastern Europe, and the other from the west (Appendix S4; S13 Fig). Further, recently sampled isolates belonging to the two populations differed in the genome-wide average *B.g. secalis* content, as per our substitution-sites approach (see above and Methods). The eastern population showed higher *B.g. secalis* proportion (18.7%) than the western population (15.4%; Welch’s t-test p < 0.0001). We then compared patterns of local ancestry along the genome in windows of 10Kb between the two groups, as obtained from MOSAIC. While estimates of mean ancestry followed similar trends between the two populations for most parts of the genome (Fig 3), two regions, one on chromosome 5 and the other on chromosome 6, stood out. Both these regions showed an excess of *B.g. secalis* ancestry in the eastern population and coincided with regions showing high divergence between the two populations (Fig. S14). These regions did not coincide with any signatures of positive selection, and we therefore cannot conclude whether these patterns emerged by chance or resulted from divergent selection in the eastern and western populations.

We also tracked how mean local ancestry changed over time by comparing isolates sampled between 2009-2013 (‘old’) with those sampled in 2022-2023 (‘new’). We could only perform this analysis for the western population because the eastern population contained no ‘old’ isolate due to under-representation of samples from Eastern Europe in the ‘old’ dataset. We found no region with drastic changes in ancestry over time (Fig. S15). Moreover, the windows associated with the highest differences in mean ancestry did not overlap with the loci putatively under selection.

## Discussion

Hybridization is a strong driver of pathogen evolution and its role in the emergence of plant diseases is widely recognized. Yet, there remain several open questions about how hybrid populations evolve and if their fate can be predicted. As genomic surveillance of plant pathogens becomes common, hybridization events can potentially be detected early, allowing their population dynamics to be tracked almost in real time. Such knowledge from a diverse range of pathogens can help bring to light the common principles of disease emergence by hybridization. Here, we present our findings on how populations of the young hybrid pathogen of triticale, *Blumeria graminis f. sp. triticale*, evolved since its emergence.

### Persistence of an isolated hybrid lineage

The earliest available genomic data for triticale powdery mildew comes from isolates sampled in 2009-2013, roughly a decade after the first reports of the disease (Mascher et al. 2006; Troch et al. 2012; Menardo et al. 2016). Population genomic analysis of these isolates showed that *B.g. triticale* formed a genetic group distinct from wheat and rye powdery mildew (Menardo et al. 2016), and our results confirmed that present day populations of this pathogen persist as the same lineage descended directly from the populations sampled previously, with no evidence of continued gene flow with the two parental *ff. spp*. The rapid establishment of a differentiated lineage, and the absence of isolates genetically intermediate between the different *ff. spp*., despite the potential niche overlap, is striking and we discuss a few hypotheses below. Importantly, we cannot rule out gene flow entirely. It is possible that *B.g. triticale* occasionally mate with one of the two parental lineages, but the eventual impact of such events on the genetic composition of the three *ff. spp*. is small enough that it could not be detected with the data and analyses presented above.

The emergence of fungal plant diseases has previously been linked to ecological speciation, and fungal life-history traits, such as production of large numbers of asexual spores and mating within hosts, have been proposed to promote rapid adaptation to new hosts and isolation from ancestral lineages (Giraud et al. 2010). In many ascomycete plant pathogens, like *Ascochyta spp*. causing blights of chickpea, and *Venturia inaequalis*, the causal agent of scab disease on apples, it has been shown that diverged populations adapted to different hosts are interfertile *in vitro* (Peever 2007; Le Cam et al. 2007), suggesting that host specificity may be the only barrier maintaining reproductive isolation. In the case of the obligate biotroph *Blumeria graminis*, it is therefore likely that host specificity contributes significantly to reproductive isolation between the *formae speciales*. While successful infections by non-adapted *ff. spp.* have been reported for many hosts including wild grasses and cereals such as wheat, rye and triticale (Troch et al. 2014; Menardo et al. 2016), these were mostly based on infection tests performed in optimal laboratory conditions on a few host varieties. In natural environments, cross-infection is probably less common, as also suggested by a previous study (Czajowski & Czembor 2016) where the authors showed that *B. graminis* isolates collected from triticale in Poland were less virulent on wheat than those collected on wheat, and vice-versa.

Our results indicate that differences in virulence spectra between wheat and triticale powdery mildew may contribute to (partial) host specificity observed in the field. In particular, the higher proportion of *B.g. tritici* isolates virulent to certain *Pm* resistance genes suggests that these genes imposed different selective pressures on the two *ff. spp*. This may be due to the lower prevalence of these resistance genes in European triticale fields, reducing selection for avirulence in *B.g. triticale* populations. A highly comparable infection assay was performed by Troch and colleagues (Troch et al. 2013) on wheat and triticale powdery mildew isolates collected in 2009 and 2010. Remarkably they found a very similar pattern, with *B.g. triticale* isolates being on average less virulent on the tested *Pm* resistance genes (see Fig. 1 in Troch et al. 2013). The only exception was *Pm8*, on which we found triticale mildew isolates to be less virulent, while Troch and colleagues reported a similar proportion of virulent isolates compared to wheat powdery mildew.

Although our results highlighted ecological separation between *B.g. tritici* and *B.g. triticale*, we did sample some cross-infections from the field and detected overlaps between their virulence profiles on different resistance genes. Furthermore, laboratory crosses between these *ff. spp*. performed *in planta*, albeit on susceptible wheat cultivars, result in viable offspring (Muller et al. 2019). It is thus possible that other post-mating barriers to gene flow, such as genomic incompatibilities, may exist between these lineages. Such barriers have been observed between sister species of other fungal pathogens, for instance in *Pyrenophora teres* (Yuzon et al. 2023), where the authors showed that hybrids between diverged lineages were less fit than the parents and identified negative epistatic interactions between parental alleles at several loci, some of which were involved in virulence. Here, we used the F1 progeny of a laboratory cross between a *B.g. tritici* isolate and a *B.g. triticale* isolate to detect regions potentially involved in such incompatibilities and found 333 inter-chromosomal pairs that did not appear to segregate independently. Though some of these regions showed high *B.g. secalis* ancestry and overlapped with signatures of divergence between *B.g. tritici* and *B.g. triticale* populations, we are cautious in interpreting these as evidence for Dobzhansky-Muller incompatibilities. A more thorough examination of these candidate regions at the population level and functional analyses of the loci involved are required to conclude if these regions indeed contribute towards isolation between the *ff. spp*. in natural populations. For example, comparing data from more *B.g. tritici* x *B.g. triticale* crosses and *B.g. tritici* x *B.g tritici* crosses can help distinguish between incompatibility pairs that are hybridization-derived and can promote isolation and those that may be involved in other epistatic interactions.

### Genome stabilization in triticale powdery mildew

When a hybrid population becomes isolated from its parental lineages, hybrid genomes proceed towards stabilization, and blocks of specific ancestry may ultimately become fixed in the population. A lack of isolation would hinder genome stabilization due to the repeated introduction of parental genomic blocks (Buerkle & Rieseberg 2008). In our study, we found that the homoploid hybrid *B.g. triticale* is established as an isolated lineage, and that between one third and one half of its genome, depending on the method used, is fixed for an ancestry in contemporary populations. Importantly, 96-98% of these fixed regions could be attributed to the *B.g. tritici* ancestry, while on average this ancestry contributes only about 71-83% to *B.g. triticale* genomes. These patterns could possibly be explained by global selection against the *B.g. secalis* ancestry due to hybridization load where mildly deleterious mutations from a parental population with a low effective population size become selected against upon entering larger populations (Juric et al. 2016; Schumer et al. 2018). The cultivation of wheat far exceeds that of rye in Europe (FAO 2025), and our efforts to sample *B.g. secalis* in 2022-2023 were largely unsuccessful, suggesting that rye powdery mildew has a smaller population size compared to wheat mildew, making selection less efficient. Moreover, loci derived from the minor parent are also expected to be selected against under models of polygenic selection if they disrupt co-adapted allele combinations and result in phenotypes far from the fitness optima (Moran et al. 2021). On average, if selection acts against hybridization, minor parent ancestry is expected to rapidly decline in the early generations, following which populations may enter a stage of slow purging (Moran et al. 2021). The average *B.g. secalis* contribution in triticale mildew appears to have remained unchanged in the last decade.

The low amount of *B.g. secalis* ancestry contributing to the *B.g. triticale* genome (17% on average) can be explained by backcrosses, selection against the ancestry from the parent with lower population size (hybrid load) (Juric et al. 2016, Schumer et al. 2018), selection against the minor parent (Moran et al. 2021), strong drift in the first hybrid generations with low population sizes, or a combination of these factors. However, a significant proportion of the genome of *B.g. triticale* appears to have stabilized within a decade after its emergence, and we would need samples from the immediate generations after hybridization to disentangle these different processes. Our finding of population substructure within *B.g. triticale*, with individuals from the eastern and western parts of Europe forming distinct groups with different amounts of *B.g. secalis* ancestry, suggest that these processes acted differently in different parts of Europe. Moreover, the initial hybridization event between wheat and rye powdery mildew was shown to have occurred at least twice independently (Menardo et al. 2016), and the differences between the eastern and western populations might be due to different origins.

Genome wide scans for signatures of recent positive selection identified several loci that were also associated with regions of fixed or nearly fixed local ancestry in contemporary populations of triticale powdery mildew. These regions were also found to be enriched for functional elements. For example, the locus on chromosome 2, which was under strong positive selection in the recent past, contains a candidate effector gene that is derived from *B.g. secalis* in all present-day *B.g. triticale* isolates. This gene is likely involved in non-host virulence. However, functional validation can be challenging as the corresponding interacting gene(s) in the host have not been characterized yet.

## Conclusion

The role of hybridization in the emergence of plant diseases is now widely recognized. As whole genome sequencing data becomes more accessible, genomic surveillance studies can track how hybrid populations evolve in multiple taxa and disentangle the underlying processes. Knowledge from different pathogens displaying distinct biological characteristics may reveal the general patterns of disease emergence through hybridization. This, in turn, can contribute to a better understanding of the dynamics of such processes and support the development of suitable control strategies.

## Materials and Methods

### Sampling and datasets

We collected 48 samples of *Blumeria graminis f. sp. triticale* over 2022 and 2023 from 32 locations spread across 8 European countries. 11 of these samples were collected from infected fields of triticale, 3 from infected fields of bread wheat and 34 from ‘trap pots’ containing young seedlings of susceptible cultivars of either wheat, rye or triticale. We also obtained 2 samples of *B.g. secalis* in 2023 – one sampled from a trap pot of a susceptible rye cultivar, and one from a collection at the Julius Kühn-Institute (JKI-A, Kleinmachnow, Germany). All 50 samples were obtained from asexual conidiospores of the fungus. In the laboratory, we propagated the isolates on 8-10 days old leaf fragments of a susceptible wheat cultivar (Kanzler) that were placed on a 0.5% agar - 0.05% benzimidazole medium in Petri dishes. Since *B.g. secalis* cannot grow on wheat, we propagated those 2 isolates on the rye cultivar Matador. For each sample, we performed two rounds of single-colony isolation to obtain pure isolates. This was achieved by first performing low-density infections on fresh, susceptible host leaves laid out on agar-containing Petri dishes. Four days after infection, the leaf fragments were observed under a binocular microscope and segments containing one colony each were cut and propagated further.

We note that it is impossible to distinguish between the different *formae speciales* of *Blumeria graminis* morphologically. We used a combination of host-specificity assays and genomic analysis to classify the individuals in our dataset (see below).

For the host-specificity assays, we used the 50 new isolates described above along with 276 *B.g. tritici* isolates sampled in 2022-2023 from Europe and the Mediterranean, as previously described in Jigisha et al. 2025. For genomic analyses, in addition to the 50 new isolates described above, we used the 625 *B.g. tritici* isolates sampled between 1980-2023 from around the world that were previously described in Jigisha et al. 2025 and passed the minimum average genome-wide coverage criterion (15x, see below). We also included publicly available genome sequences of other closely related lineages: 2 samples of *B.g. dactylidis*, 22 *B.g. triticale*, 5 *B.g secalis* and 7 *B.g. dicocci* isolates that had been collected between 1990-2013 from Europe and the Middle East. The metadata associated with all samples, including their accession numbers, are listed in Table S1.

### Host specificity assays

We tested the host specificity of the 50 new isolates of *Blumeria graminis* sampled in this study along with 275 (of 276) *B.g. tritici* isolates collected in Jigisha et al. 2025. One *B.g. tritici* isolate described in Jigisha et al. 2025 was not phenotyped (ITCN052201). The array of hosts tested included hexaploid wheat cultivar Kanzler, two triticale cultivars - Bedretto and the Agroscope breeding line 1011.58329 (hereon referred to as “Agroscope”), and a rye cultivar Matador. Ten days old leaf fragments placed in agar plates were infected with conidiospores and incubated at 20C under a 16h light 8h dark cycle. We used three biological replicates for each cultivar. Images of the infected leaves were taken ten days post infection. All isolates that were able to successfully infect Kanzler, Bedretto and Agroscope were classified as *B.g. triticale*.

### DNA extraction and whole-genome sequencing

We extracted DNA for all isolates following a protocol using magnetic beads (Rohland & Reich 2012) adapted for the KingFisher Apex 96 System. Short-read, paired-end (150 bp, insert sizes ca. 200-350 bp) genome sequencing was performed for all isolates using Illumina Truseq Nano libraries and the Illumina NovaSeq 6000 or NovaSeq X Plus instruments.

### Variant Calling

Raw, paired-end reads from all 50 newly sequenced isolates and 36 downloaded sequences of *B.g. secalis*, *B.g. triticale*, *B.g. dicocci* and *B. dactylidis* (S1 Data) were filtered and mapped to a reference genome for subsequent variant calling following the same steps used in Jigisha et al. 2025. Briefly, fastp v0.23.2 (Chen et al. 2018; Chen 2023) default settings were used to trim adapters and quality-based trimming was executed using – cut_front (window size 1, mean quality 20) and –cut_right (window size 5 and mean quality 20). fastp –merge was used to merge overlapping paired-end reads, with overlap_len_require = 15 and overlap_diff_percent_limit = 10. All reads were mapped to the updated 96224 *B.g. tritici* reference genome (Müller et al. 2019) which included 11 chromosomes, the mitochondrial genome (Zaccaron & Stergiopoulos 2021), a contig of the alternate mating type and an ‘unknown chromosome’ with scaffolds that could not be assigned to any chromosome (Jigisha et al. 2025). The alignments were sorted and merged, and genome wide coverage was calculated with Samtools v1.17 (Danecek et al. 2021). All samples had a genome-wide coverage of 15x or more. Each sample was assigned one of two mating types based on the coverage over the mating type genes. GATK v4.4.0.0 HaplotypeCaller (Van der Auwera et al. 2020) was then used for sample-level haplotype calling.

For variant calling, we combined the 86 individual VCF files with the per-sample VCF files of 625 *B.g. tritici* isolates generated in Jigisha et al. 2025. *The B.g. tritici* isolates that were found to have less than 15x coverage in Jigisha et al al. 2025 were not included in this study. All per-sample VCF files were merged using GATK CombineGVCFs to perform joint genotyping with GATK GenotypeGVCFs. We used GATK VariantFiltration to exclude low-quality sites using the filters QD<10, FS>55, MQ<45 and –4<ReadPosRankSum<4. Additionally, for every site, we recoded calls with < 90% read support (heterozygous calls) or those with depth of informative, high-quality reads less than 8 (DP < 8) as missing data. Following Jigisha et al. 2025, we identified potential mixed infection by looking for samples for which the ratio of the number of variants to the number of heterozygous sites was lower than one. We identified 14 such samples (12 *B.g. tritici* and 2 *B. dactylidis)* and excluded them from all analyses. Additionally, we also excluded from further analyses the 45 *B.g. tritici* isolates that were identified as suspected contaminants and clones in Jigisha et al 2025. In the final VCF file, 652 isolates were retained.

### Genetic clustering of *formae speciales*

Though the *formae speciales* classification is based on phenotypes, i.e., which hosts a pathogen can infect, it has been shown previously that in *B. graminis* the different *formae speciales* also form distinct groups genetically. We performed a principal component analysis (PCA) to see how the different samples clustered based on their genomes. We included all 652 isolates and filtered out singletons and sites with more than 10% missing data using GATK SelectVariants to obtain 2,431,821 biallelic SNPs over the 11 chromosomes. The PCA was performed using the function glpca in the R package adegenet (Jombart & Bateman 2008; Jombart & Ahmed 2011) and the results were visualized using ggplot2 (Wickham 2016).

### Identification of clones

We identified clonal isolates of *B.g. triticale* based on the number of SNPs between pairs of individuals. We calculated a pairwise distance matrix for all pairs of individuals with the dist.gene function of the R package ape (Paradis & Schliep 2019). Upon evaluating the distribution of the distances, we classified all pairs of isolates with less than 9e-05 nucleotide differences per site between them as clones of each other. There were 5 clonal pairs and 1 group of 4 clonal isolates. Each group contained isolates collected from the same location and we retained only one isolate from each group for all subsequent analyses. In other words, we excluded 8 (of 70) *B.g. triticale* isolates from all downstream analyses.

### Phylogenetic network, population structure and population tree

We constructed a phylogenetic network for 415 *B.g tritici* isolates, 62 *B.g. triticale* isolates and 7 *B.g. secalis* isolates sampled from Europe and neighbouring regions (total n = 484). We used SplitsTree App 6.0.0 (Huson & Bryant 2024), with the pairwise genetic distance matrix described above as input. The Neighbour Net method (Bryant & Moulton 2004; Bryant & Huson 2023) was used with default options resulting in 526 splits. The Show Splits method was used to visualize the Split Network.

We used fineSTRUCTURE (Lawson et al. 2012) to characterize the fine-scale structure in our data while making use of linkage information. We used the 484 individuals described above and retained 1,293,329 biallelic SNPs over 11 chromosomes with no missing data. The genetic map used to calculate the per-base recombination rates has been described previously (Müller et al. 2019). The chromopainter steps, as executed in stages 1 and 2 of fineSTRUCTURE, were performed with default settings. The inferred values of mu, Ne and c were 0.001153, 72.3377 and 0.3959 respectively. For the MCMC inference step, we used the greedy optimization option with default settings which achieved approximate convergence after 4 iterations.

Based on the fineSTRUCTURE results, we divided the set of 484 isolates into 8 groups, namely – *B.g. secalis*, *B.g. triticale* (EAST), *B.g. triticale* (WEST), *B.g. tritici* (N_EUR, S_EUR1, S_EUR2, TUR and ME). We used TreeMix v 1.13 (Pickrell & Pritchard 2012) to construct a population tree while accommodating gene flow by introducing migration edges. We included 799,814 biallelic SNPs from chromosomes 1-11 after removing all sites with missing data and singletons. To account for linkage disequilibrium, we grouped together SNPs in blocks of size 1000 using the –k option in TreeMix. We sequentially added migration edges, going from 0 to 10, and visualized the proportion of variance explained, residuals, and the resulting tree for each model.

### Quantifying parental genomic contribution

We quantified the contribution of *B.g. secalis* and *B.g. tritici* in the genomes of all *B.g. triticale* (n = 62) individuals and in *B.g. tritici* sampled in 2022 and 2023 (n = 255; *Europe+_2022_2023* dataset from Jigisha et al. 2025) using an approach was similar to that described in (Menardo et al. 2016; Müller et al. 2021). This method did not make use of any formal probabilistic model and relied instead on a set of diagnostic SNPs that were fixed for different alleles in the two “donor populations.” We refer to these sites in the genome as “substitution sites”. We used two donor panels, one comprising all sampled *B.g. secalis* genomes (n = 7) and the second comprising *B.g. tritici* individuals (n = 17) that had been sampled from Europe in or before 2001, the year of emergence of powdery mildew on triticale. Using these donor panels, we identified 361,578 diagnostic SNPs or substitution sites spread across 11 chromosomes. Next, we characterized which donor allele was present at these sites in each of the 317 target individuals. Based on this, we assigned ancestry to continuous stretches of the target genomes, extending segments if consecutive sites were found to be from the same donor. If the distance between two sites was greater than 10 Kb, we considered this region to be uninformative or ‘missing data’. Blocks with length 1 were also considered missing data.

### Phylogenetic analysis of mating type genes

The reference isolate 96224 has a MAT1-2 mating type, and two genes associated with this mating type are annotated on chromosome 1 (*Bgt_2805* or *SLA2*, and *Bgt_3306* or *MAT1-2-1;* Müller et al. 2019). The alternative mating type is MAT1-1, and in the genome assembly used in this study, one contig containing a part of the MAT1-1 locus was included, so that all sequenced isolates have been mapped on both mating types. The MAT1-1 locus, however, was not annotated. We used STAR v. 2.7.11b with default parameters (Dobin et al. 2013) to map RNAseq reads for isolate 94202 (Praz et al. 2018) (GEO accession number GSE108405), and we used IGV v. 2.16.2 (Thorvaldsdóttir et al. 2013) to visualise the read alignments on the MAT1-1 locus and annotate the only gene present. The annotation resulted in a 1,008-nucleotide gene with 100% homology to the partial sequence of the Bgt alpha box mating-type protein MAT1-1-1 gene on NCBI (accession: HQ171902.1).

For all three genes, we performed a phylogenetic analysis based on their genomic sequence. We extracted the nucleotide sequences of all isolates from the first position of the coding sequence to the last (excluding the stop codon). These sequences included intronic sequences. All indels were recoded as missing data. The pseudo-alignments composed of invariant sites and SNPs were used for a phylogenetic analysis with raxml-ng (Kozlov et al. 2019) with model GTR + G.

### Infection tests and virulence analysis

To test for differences in the ecological niches of *B.g. tritici* and *B.g. triticale* we performed infection assays on 20 wheat lines carrying either single *Pm* genes or a combination of multiple Pm genes. We tested 34 *B.g. triticale* isolates and 104 *B.g. tritici* isolates collected in 2022 and 2023 from France, Germany, Netherlands, Poland and Hungary (Table S1). Each isolate was tested on at least five plants for each line, as well as Nimbus plants as the susceptible control for inoculation effectiveness. Seedlings were inoculated at approximately 10 days old with fully expanded first leaves (DC: 12) (Zadoks et al. 1974). After inoculation, plants were grown at 19°C/15°C day/night temperature and a 16-hour photoperiod. Host reactions were scored after approximately 8-10 days, once the fungal mycelium was fully developed on the susceptible check. Infection types were indicated according to a 5-level scale (Mains & Diktz 1930) where 0, 1, and 2 represented resistant plants (0 means immune, i.e., no visible infection symptoms; 1 – hypersensitive reaction with necrotic flecks; 2 - small colonies with necrotic flecks, no or scarce sporulation) and 3 and 4 represented susceptible plants (3 - moderate mycelial growth and sporulation, small necrotic areas; 4 - well-developed mycelium and good sporulation). To prevent mildew contamination, the plants infected by various isolates were grown separately in transparent boxes. Differences in virulence among isolates were visualized with a categorical PCA with each variable considered ordinal. The analysis was performed with the princal function from the R package Gifi (Mair et al. 2022).

### Dobzhansky-Muller genetic incompatibilities

To test for genomic incompatibilities between *B.g. tritici* and *B.g. triticale*, we made use of the mapping population generated in (Müller et al. 2019) where the authors crossed one *B.g. tritici* isolate (96224) with one *B.g. triticale* isolate (THUN-12) in the laboratory and sequenced the resulting offspring. The paired-end, short-read, whole-genome sequences of the 118 progenies are publicly available under the accession number SRP148738. We downloaded these sequences using fasterq-dump from the SRA Toolkit (https://hpc.nih.gov/apps/sratoolkit.html) with the ‘split-files’ option. The raw reads were filtered and aligned to the 96224 *B.g. tritici* reference genome using the same steps as described above (see section ‘Variant calling’). We excluded 2 progenies with average genome-wide coverage of less than 5x. The sample-level VCF files of the remaining 116 progenies, as generated by GATK v 4.4.0.0 HaplotypeCaller, were combined with those of 96224 and THUN-12 (as generated earlier) using GATK CombineGVCFs for subsequent joint-genotyping with GATK GenotypeGVCFs. Hard filtering for sites was performed using GATK Variant Filtration, excluding sites with QD < 10, FS > 50, MQ < 45 and ReadPosRankSum beyond the range (–5, 5). Sites with spanning deletions were also excluded. Additionally, genotype calls with < 90% read support and/or read-depth less than 5 were recoded as missing data.

We used the genetic map from (Müller et al. 2019) to identify non-recombinant windows (i.e. genomic segments in which no recombination event was observed in the 96224xTHUN-12 cross). We found 2604 such windows and for each progeny we determined the parent of origin using the 257,021 chromosomal SNPs between 96224 and THUN-12 (diagnostic SNPs). We excluded windows with five or less diagnostic SNPs. For each progeny, if more than 50% of diagnostic SNPs within a window had missing genotypes, or if fewer than 99.9% of the non-missing diagnostic SNPs matched the genotype of one single parent, the window was considered unassigned (i.e., missing data).

We considered all possible pairwise combinations between windows and assembled a matrix containing the combinations of parental genotypes for each of the 116 progenies. For each pairwise window comparison we calculated the odd ratio (OR) as (a*d)/(b*c) where a, b, c, and d, are respectively the number of progenies with a 96224/96224, 96224/THUN-12, TUHN12/96224, and THUN-12/THUN-12 genotype. We tested whether the OR was significantly different from 1 with a Fisher exact test. We identified 334 windows with OR > 1 and false discovery rate (Benjamini &Hochberg 1995) corrected p-value < 0.01. For these windows we performed 100 permutations in which we reassigned the parental origin at random among progenies (within each window). We then used these permuted datasets to calculate the odd ratio. We considered as significant only comparisons with a false discover rate (Benjamini-Hochberg) corrected p-value < 0.01, and with a higher OR compared to the maximum OR obtained with the permuted datasets (empirical p-value < 0.01).

We computed measures of population divergence (d_xy_ and F_ST_) between *B.g. triticale* (n = 62) and the *B.g. tritici* N_EUR population (n = 212) in non-overlapping windows of 10 Kb long chromosomes 1-11 using pixy (Korunes & Samuk 2021). For visualization and comparison with putative DMIs, we extracted genome windows with the highest 10% of each of the summary statistic values. The incompatible pairs, parental genomic contribution in THUN-12 and the d_xy_, F_ST_ outliers were visualized together in a circle plot using the R package BioCircos (Vuillard et al. 2019).

### Local ancestry estimation and genome stability analysis

We took a probabilistic approach to estimate local ancestry along *B.g. triticale* genomes using the R package MOSAIC v 1.5.0 (Salter-Townshend & Myers 2019). This method uses nested Hidden Markov Models (HMM) and does not require that the donor panels accurately represent the admixing groups. Instead, it infers the relationship between the ‘ancestral’ admixing groups and the donor panel from the data. We first ran MOSAIC with the same donor panels used to quantify genome contribution based on substitution sites (17 *B.g. tritici* and 7 *B.g. secalis* individuals; see above). Additionally, in another run, we replaced the old *B.g. tritici* donor panel with a new one, comprising 175 *B.g. tritici* individuals that were sampled in 2022-2023 and belonged to the N_EUR population, as inferred from fineSTRUCTURE. We filtered out all sites with any missing data, as well as sites present in ambiguous regions on the genetic map, which resulted in 1,088,697 SNPs over 11 chromosomes across 261 individuals. MOSAIC was run to model two-way admixture, independently for each set of donor panels, with the parameters –nophase and –Ne 1290 (approximated as θ/2µ, where θ is Watterson’s theta estimated using the number of segregating sites and the number of individuals, and µ is the mutation rate, taken to be 5 x 10^-7^ mutations per base pair per generation following Sotiropoulos et al. 2022). The local ancestry estimates, originally computed for evenly spaced positions on genetic distance, were translated to estimates on physical SNP positions using built-in functions of MOSAIC. The two ancestries inferred in both runs of MOSAIC were well represented by the donor panels.

We then assessed genome stability in *B.g. triticale* by characterizing variation in local ancestry along the genome among individuals sampled from different geographical locations and at different time periods. We used the local ancestry estimates of the *B.g. secalis*-dominant ancestry at SNP positions along the genome, as obtained from the two independent runs of MOSAIC to calculate mean *B.g. secalis* ancestry in non-overlapping windows of 10 Kb across chromosomes 1-11. We obtained 9919 such windows, each with at least one SNP, covering 72.2% of the whole genome. For each isolate, windows with mean ancestry greater than 0.7 were assigned *B.g. secalis* ancestry, and those with mean ancestry less than 0.3 were assigned *B.g. tritici* ancestry. Windows that were assigned to the same ancestry across both runs of MOSAIC in all members of a population were considered fixed for that ancestry in that population. Population-level mean ancestry was also obtained in windows of 10 Kb by averaging over the mean ancestry across all individuals of a population for each window.

We also compared the results obtained from the two iterations of MOSAIC to the substitution-sites approach. For the sake of comparison, we divided the genomes of *B.g. triticale* isolates into windows of 10 Kb and obtained 10685 such windows (covering ∼ 77.8% of the genome) with least one SNP or substitution site that could be assigned to a ‘parent’ across all isolates. Each SNP was assigned a value of 0 if it was of *B.g. tritici* origin, and 1 if it was of *B.g. secalis* origin. We then computed the mean for each window and for each isolate. If the window had only *B.g. secalis* SNPs (i.e. mean = 1) across all members of a population, then that window was said to be fixed for *B.g. secalis* ancestry in that population, and if the window-mean was 0 across all isolates, then that window was fixed for *B.g. tritici* ancestry.

### Population structure within *B.g. triticale*

We characterized the distribution of genetic variation within *B.g. triticale* using different methods. We performed a PCA for the 62 non-clonal *B.g. triticale* individuals using 536,386 biallelic SNPs spread over chromosomes 1-11. We filtered out all singletons and sites with more than 10% missing data. The PCA was performed using the glpca function from the R package adegenet (Jombart & Ahmed 2011; Jombart & Bateman 2008). Next, to run the ADMIXTURE analysis (Alexander et al. 2009), we performed LD based pruning of the set of SNPs used in the PCA using PLINK v 1.9 (Chang et al. 2015) (–indep-pairwise, window size = 25kb, step size = 1 SNP, correlation threshold r2 = 1). This resulted in 6,297 biallelic SNPs over 11 chromosomes. We ran 10 replicates of ADMIXTURE for each K from 1 to 10 and visualized the cross-validation error across all runs to infer the best value of K(K=2). We also performed the fineSTRUCTURE analysis (Lawson et al. 2012) for *B.g. triticale*. We filtered out all sites with any missing data, retaining 521,983 biallelic SNPs over 11 chromosomes. The per-base recombination rates were calculated using the genetic map from Müller et al. 2019, as described in Jigisha et al. 2025. FineSTRUCTURE was run with the default settings with ploidy 1, and the estimated values of Ne and c were 1984.38 and 1.06483 respectively.

For between-population comparisons, we divided our dataset into 2 populations – east and west, each comprising individuals that were consistently grouped together in the different runs of fineSTRUCTURE and other population structure analyses. 7 individuals were excluded on grounds of ambiguous population assignment. We computed d_xy_ and F_ST_ between the eastern and western populations in windows of 10 Kb along the genome using pixy (Korunes & Samuk, 2021).

### Genome-wide scans for selection

We performed genome-wide scans for selection using isoRelate (Henden et al. 2018), a method that detects signatures of recent positive selection by identifying regions of the genome that show excess identity-by-descent in a population. We ran this analysis for all *B.g. triticale* individuals together, using biallelic SNPs. We excluded sites with any missing data, those with minor allele frequence less than 0.05 and those that could not be mapped unambiguously on the genetic map. IsoRelate was run to identify identical-by-descent segments that were larger than 2cM and 50 Kb, and that had at least 25 SNPs.

We used bedtools intersect (Quinlan & Hall 2010) and the publicly available GFF file associated with the 96224 genome (Müller et al. 2019) to identify genes that overlapped with the signatures of selection. The list of known fungicide targets and avirulence genes was obtained from published resources (Bernasconi et al. 2025 a&b; Bourras et al. 2015; Bourras et al. 2019; Hewitt et al. 2021; Kloppe et al. 2023; Kunz et al. 2023; Kunz et al. 2025; Minadakis et al. 2025; Müller et al. 2022; Praz et al. 2017). The loci found to be under selection in *B.g. triticale* were also compared with those found in the five *B.g. tritici* populations described in Jigisha et al. 2025.

### Genome alignment and haplotype network

The 96224 and THUN-12 reference assemblies (GenBank IDs GCA_900519115.1 and GCA_905067625.1 respectively) were aligned using the nucmer script from MUMmer v4.0.1. To focus on the the region of interest on chromosome 2, the output delta file was parsed through the dnadiff script of MUMmer and the resulting .1coords file was used to generate the synteny plot using the R package gggenomes (Hackl et al. 2024). The gene annotations for both genomes were taken from the publicly available GFF files.

The nucleotide sequence corresponding to *BgtE-5850* (chromosome 2: 9305082-9305589) was extracted from the all-sites VCF file for all 484 *B. graminis* isolates in our dataset (415 *B.g. tritici*, 62 *B.g. triticale*, 7 *B.g. secalis*). This was used to construct a haplotype network using the haploNet function implemented in the R package pegas (Paradis et al. 2010).

## Data Availability

The short read genome sequence data generated in this study is available under the BioProject Accession PRJEB96281. Accession numbers for all isolates used in this study are listed in Table S1. The code to reproduce the analyses presented here is available at https://github.com/jjigisha/bg_triticale. All other data are contained within the manuscript and its supplementary information.

## Acknowledgements

We thank Beat Keller, Gerhard Herren, Helen Zbinden, Dolors Villegas, Kerstin Flath and Susan Bergmann for their support. We also thank all colleagues who contributed to sampling powdery mildew. Part of the genome sequencing was performed at the Functional Genomics Center Zurich (FGCZ) of the University of Zurich and ETH Zurich. All computation work was performed on the ScienceCluster provided by the Science IT team of the University of Zurich. This work was funded by the Swiss National Science Foundation grant number PZ00P3_193473 awarded to FM.

