## Supplementary Information for "Tracking the evolutionary trajectory of a young hybrid plant pathogen"

### Contents

**1 Figures** **1**

**2 Appendices** **15**

**References** **18**

### Figures

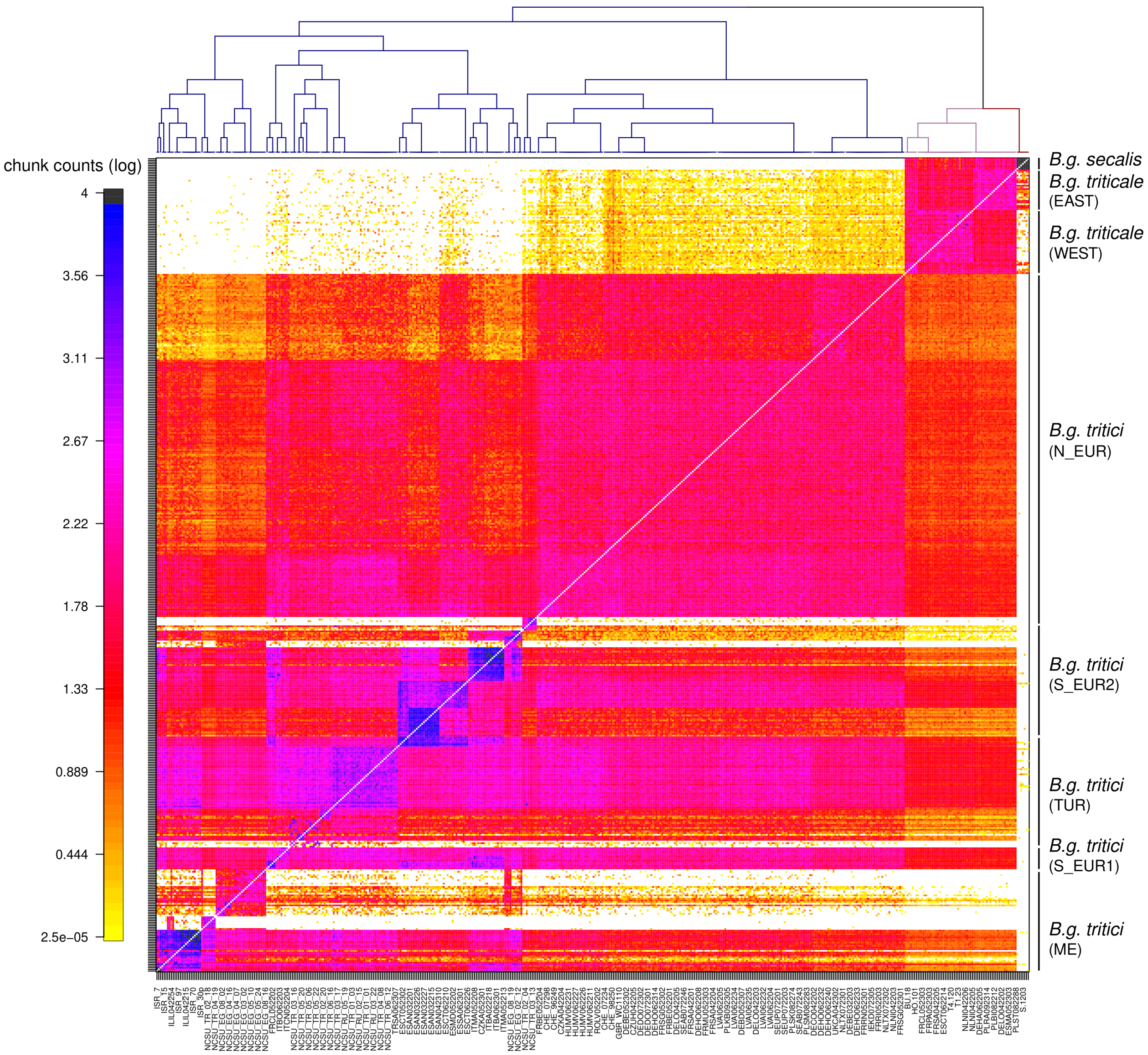


**Fig. S1. fineSTRUCTURE analysis of *Blumeria graminis***
Dendrogram (on top) and coancestry matrix computed by fineSTRUCTURE for 484 *Blumeria graminis* isolates (*ff. spp. secalis, triticale* and *tritici*) sampled from Europe and neighbouring regions. Colours in the matrix represent chunk count in log scale, which is the number of genomic segments donated by isolates in rows to isolates in columns. Higher values of chunk count reflect higher shared ancestry.

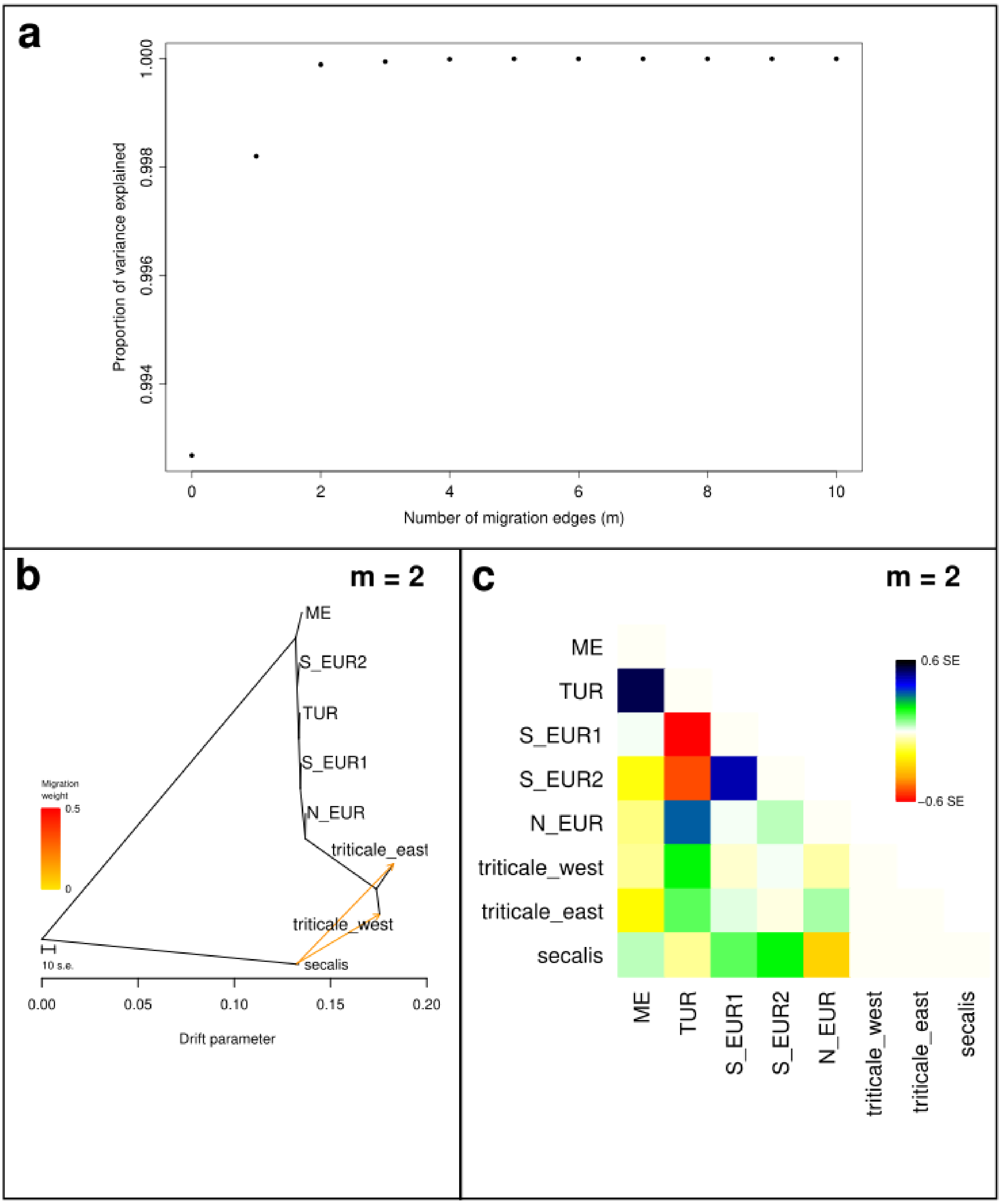


**Fig. S2. TreeMix analysis of *Blumeria graminis***
(**a**) Proportion of variance explained by each independent run of TreeMix with different numbers of migration edges. (**b**) Population tree and (**c**) residuals for the TreeMix run with 2 migration edges. ME, S_EUR2, TUR, S_EUR1 and N_EUR are *B.g. tritici* populations, triticale_east and triticale_west are the subpopulations of *B.g. triticale* and secalis is comprised of all *B.g. secalis* isolates. Positive values of residuals indicate that adding an additional migration edge may improve the fit of the model, while the reverse is true for negative residuals.

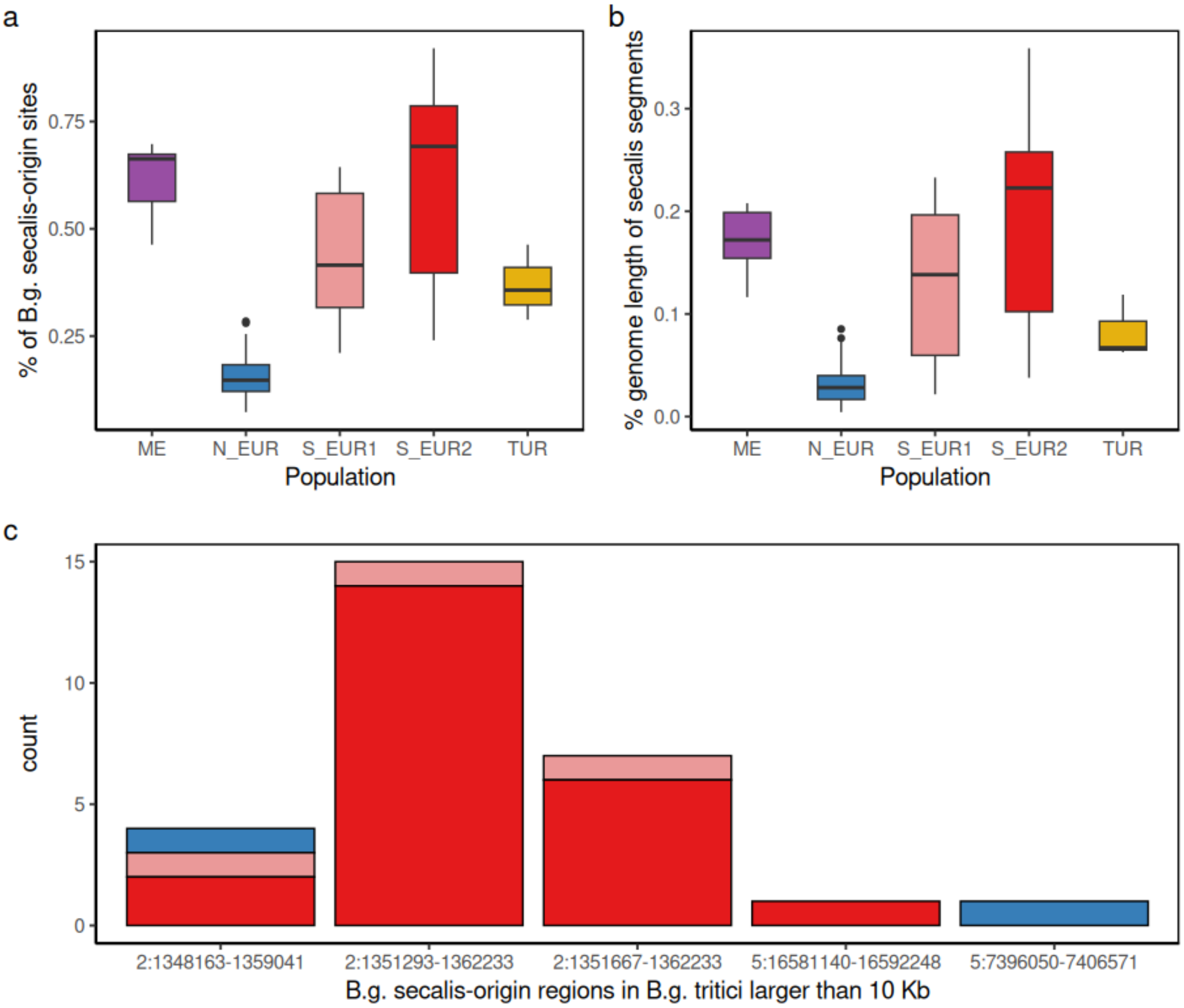


**Fig. S3. *B.g. secalis* origin regions in new *B.g. tritici* isolates**
(**a**) Genome-wide percentage of substitution-sites that could be assigned to *B.g. secalis*-origin in *B.g. tritici* isolates sampled in 2022-2023, subdivided by population. (**b**) Percentage of the genome covered by *B.g. secalis* origin segments (**c**) Frequency of occurrence of *B.g. secalis*-origin segments with length greater than 10 Kb. Labels on x axis represent genomic coordinates (chr:from bp - to bp).

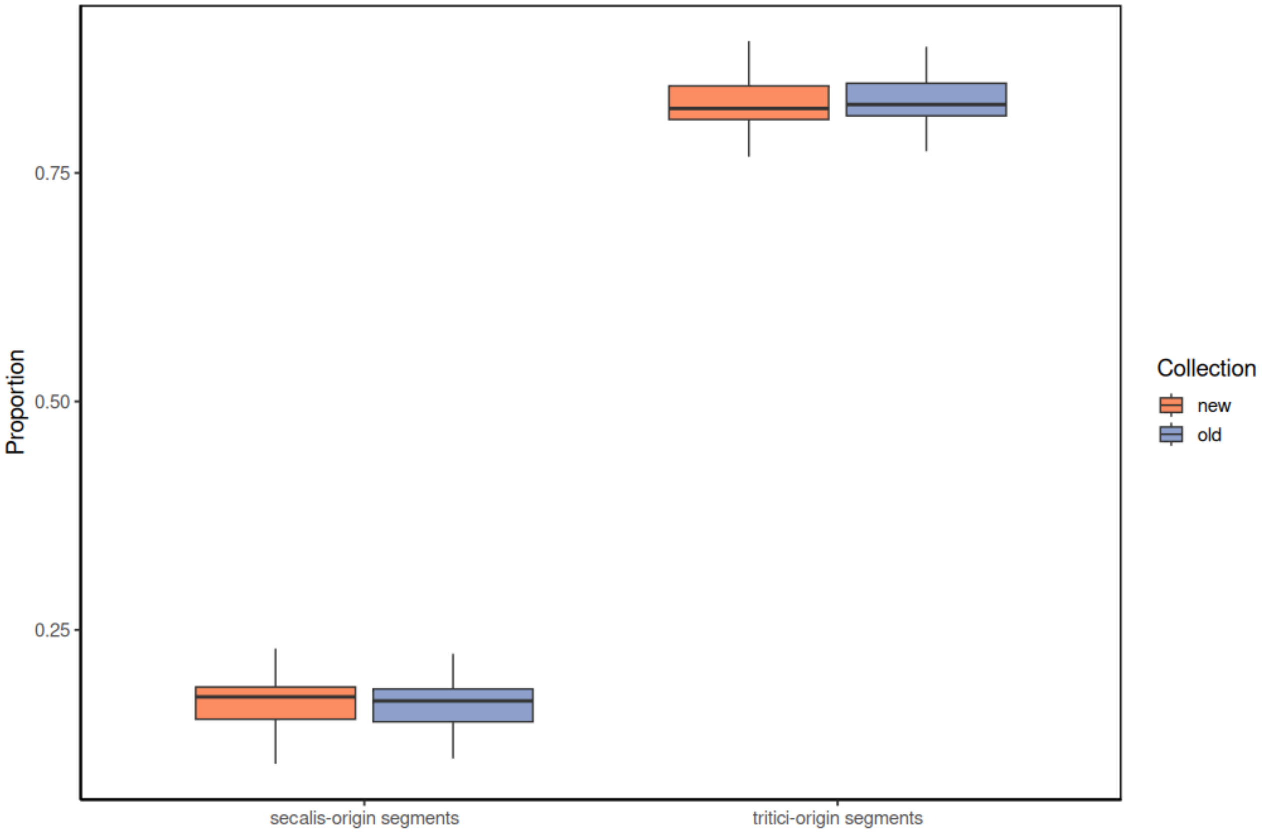


**Fig. S4. Differences in parental contribution between old and recently sampled *B.g. triticale***
The proportion of the genome covered by *B.g. secalis* origin segments and *B.g. tritici* origin segments, as inferred using the substitution sites approach (see Methods). The 'new' collection comprises 46 *B.g. triticale* isolates sampled in 2022-2023 and the 'old' collection contains 16 *B.g. triticale* isolates sampled between 2009-2013.

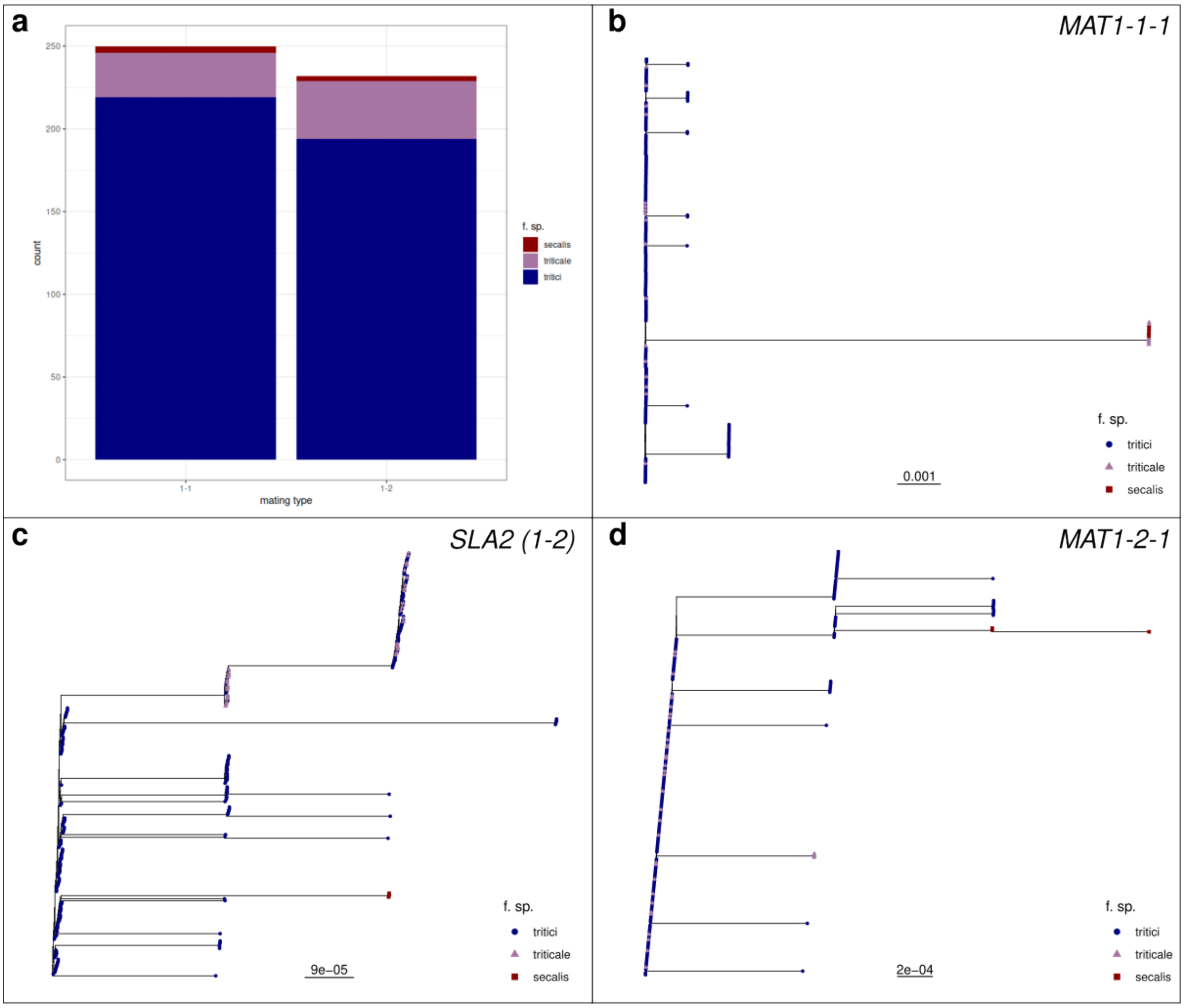


**Fig. S5. Phylogenetic analysis of mating type genes**
(**a**) Frequency of occurrence of the two mating types, 1-1 and 1-2 in the three *ff. spp.* (**b**) Phylogenetic tree of the *MAT1-1-1* gene associated with the 1-1 mating type, and (**c**) *SLA2* and (**d**) *MAT1-2-1* genes associated with the 1-2 mating type. The colours represent the *ff. spp.* and are consistent across all panels.

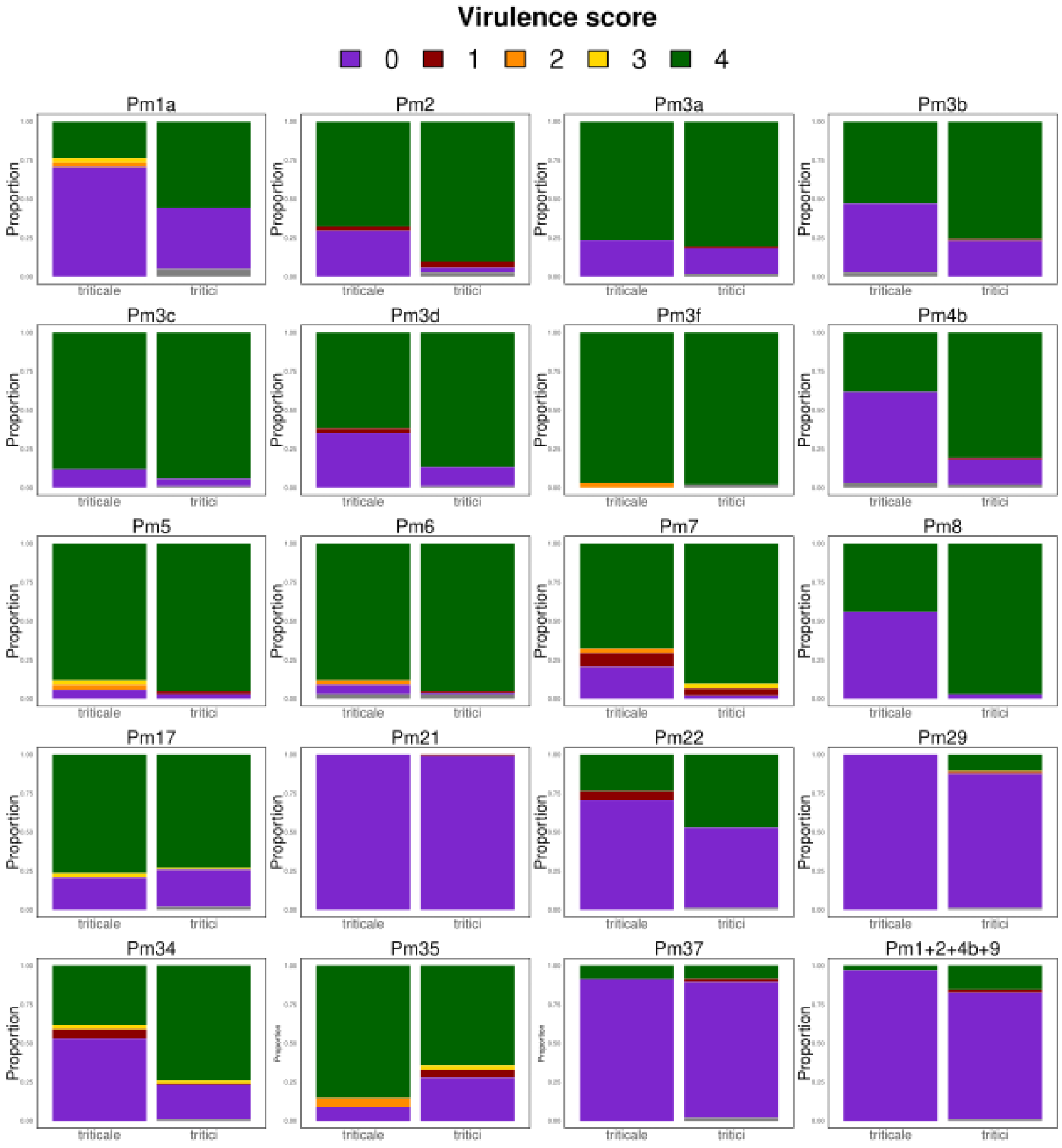


**Fig. S6. Results from infection assays on wheat lines containing different Pm genes**
Each panel shows the results from the infection assay on a different wheat line. The title for each panel indicate the resistance (Pm) gene(s) carried by each line. The bars show the proportion of *B.g. triticale* ('triticale') and *B.g. tritici* ('tritici') isolates that were assigned the different virulence scores. A virulence score of 0 means the isolate was avirulent, i.e. no visible symptoms of infection. Scores from 1 to 4 reflect different host responses and levels of mycelial growth, with 4 being the most virulent (see Methods).

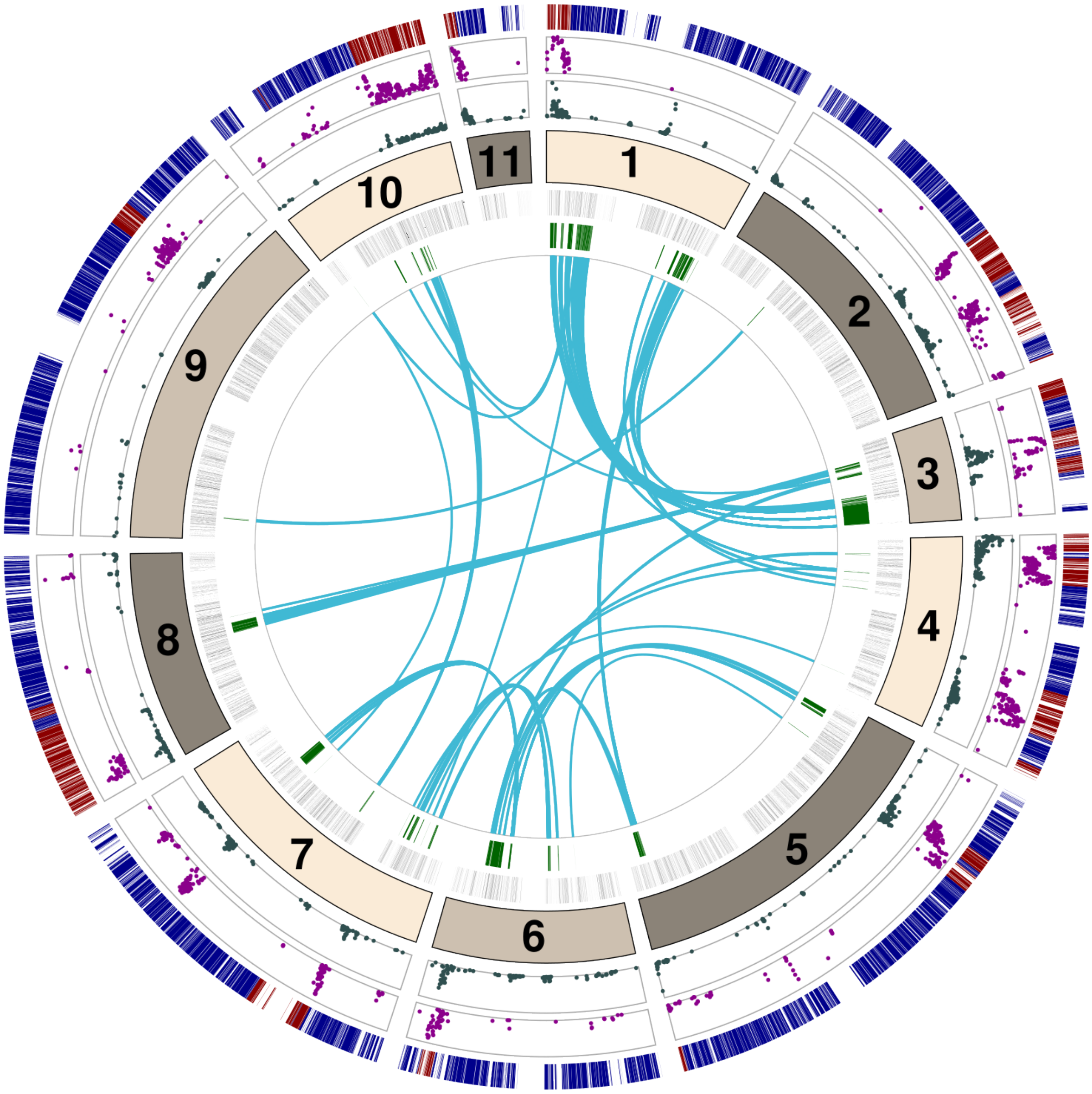


**Fig. S7. Candidate Dobzhansky-Muller incompatibilities between wheat and triticale powdery mildew**

The numbered track represents the 11 chromosomes of *Blumeria graminis*, with sizes proportional to the length of each chromosome. Pairs of genomic regions that did not segregate independently in the F1 offspring of 96224 x THUN-12 were identified as candidate DMIs. These regions are marked in bright green in the inner most circle and each blue arc represents one of the 333 DMI pairs. The grey segments represent all the annotated genes in the 96224 reference assembly. The dark green and magenta points indicate the value of $d_{xy}$ and $F_{ST}$ estimates respectively, obtained upon comparing *B.g. triticale* and the *N_EUR* population of *B.g. tritici*. Only the genomic windows within the top 10% for each statistic were plotted. The outermost circle shows the genome segments that could be assigned as *B.g. tritici*-origin (blue) or *B.g. secalis*-origin (red) in THUN-12 using the substitution-sites method.

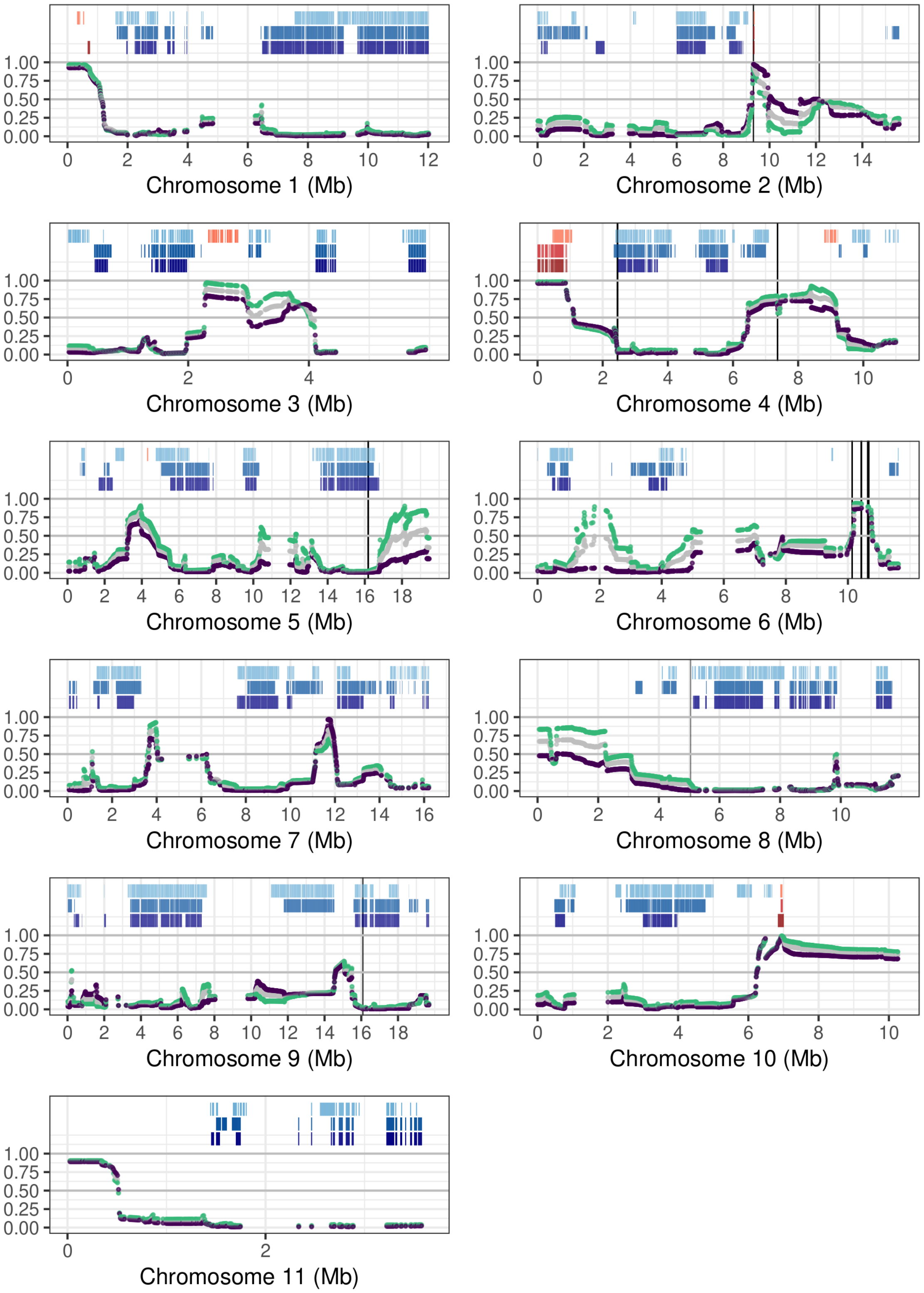


**Fig. S8. Variation in local ancestry in contemporary *B.g. triticale* populations**
The blue and red segments at the top of each panel (chromosome) represent the genomic windows fixed for *B.g. tritici* or *B.g. secalis* ancestry respectively across all *B.g. triticale* isolates sampled in 2022-2023. The first row from the top shows results from the substitution sites method, the second row corresponds to the MOSAIC run with old *B.g. tritici* donors and the third belongs to the MOSAIC run with new *B.g. tritici* donors. Plotted in the lower section of each panel is mean *B.g. secalis* ancestry in windows of 10 Kb, as obtained from the MOSAIC run with new donors. Mean ancestry across all *B.g. triticale* isolates sampled in 2022-2023 is shown in grey, while green and purple correspond to mean ancestry for the eastern and western populations respectively. The vertical black lines represent loci found to be under recent positive selection from the isoRelate analysis, after FDR correction.

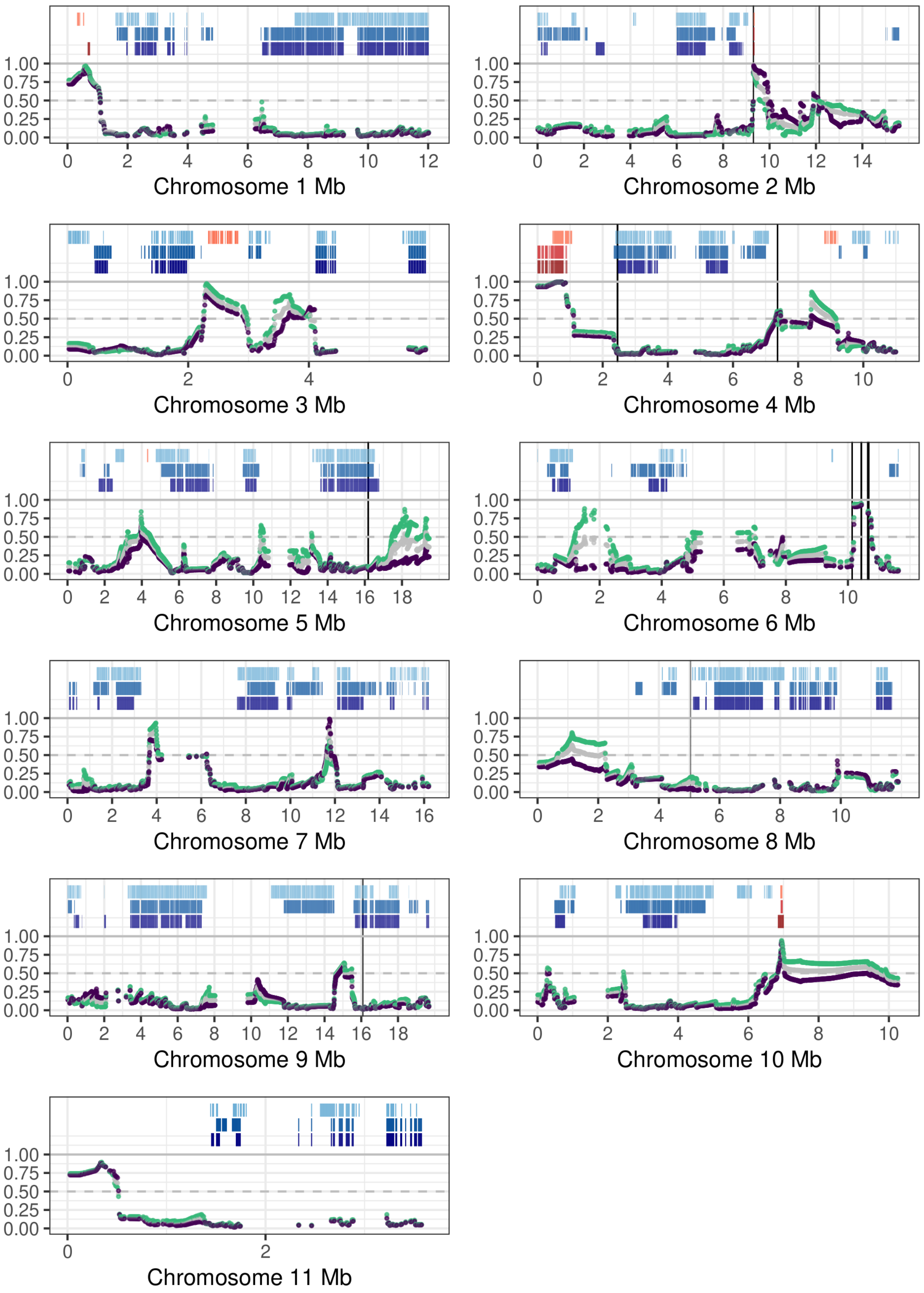


**Fig. S9. Variation in local ancestry in contemporary *B.g. triticale* populations**
This figure is the same as Fig. S8, except that the mean *B.g. secalis* ancestry estimates plotted here were obtained from the MOSAIC run with the old *B.g. tritici* donors, instead of the iteration with the new *B.g. tritici* donor panel.

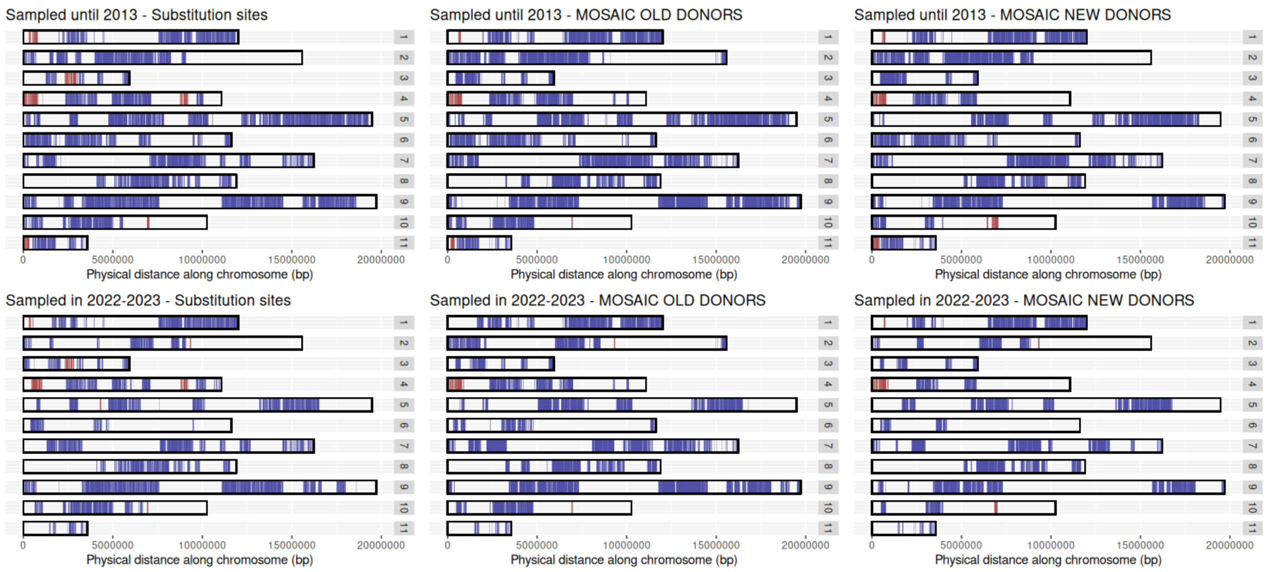


**Fig. S10. Comparing regions fixed for an ancestry in old versus new *B.g. triticale* isolates**
Each panel shows the 11 chromosomes. Red represents the 10 Kb windows that show *B.g. secalis* ancestry across all *B.g. triticale* isolates sampled until 2013 (top 3 panels) and across all *B.g. triticale* isolates sampled in 2022-2023 (bottom 3 panels). Substitution sites, MOSAIC OLD DONORS and MOSAIC NEW DONORS are the three methods we used to estimate local ancestry.

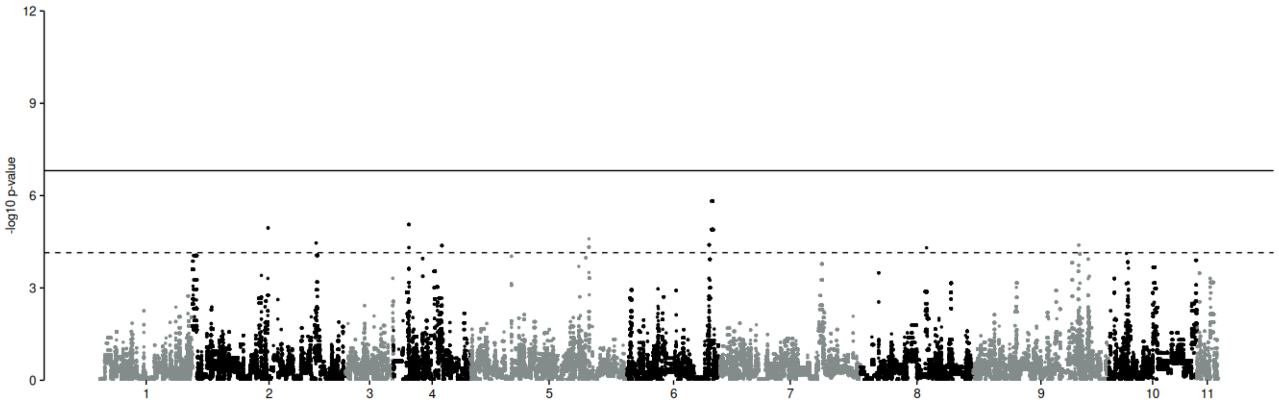


**Fig. S11. Genome-wide scan for signatures of recent selection**
Manhattan plot produced by the isoRelate analysis of *B.g. triticale.* The y-axis represents the significance of the excess of sharing identical-by-descent segments between pairs of isolates. The dashed horizontal line shows the False Discovery Rate (FDR) corrected 0.05 threshold while the full horizontal line shows the Bonferroni corrected 0.05 threshold (in -$\log_{10}$ scale).

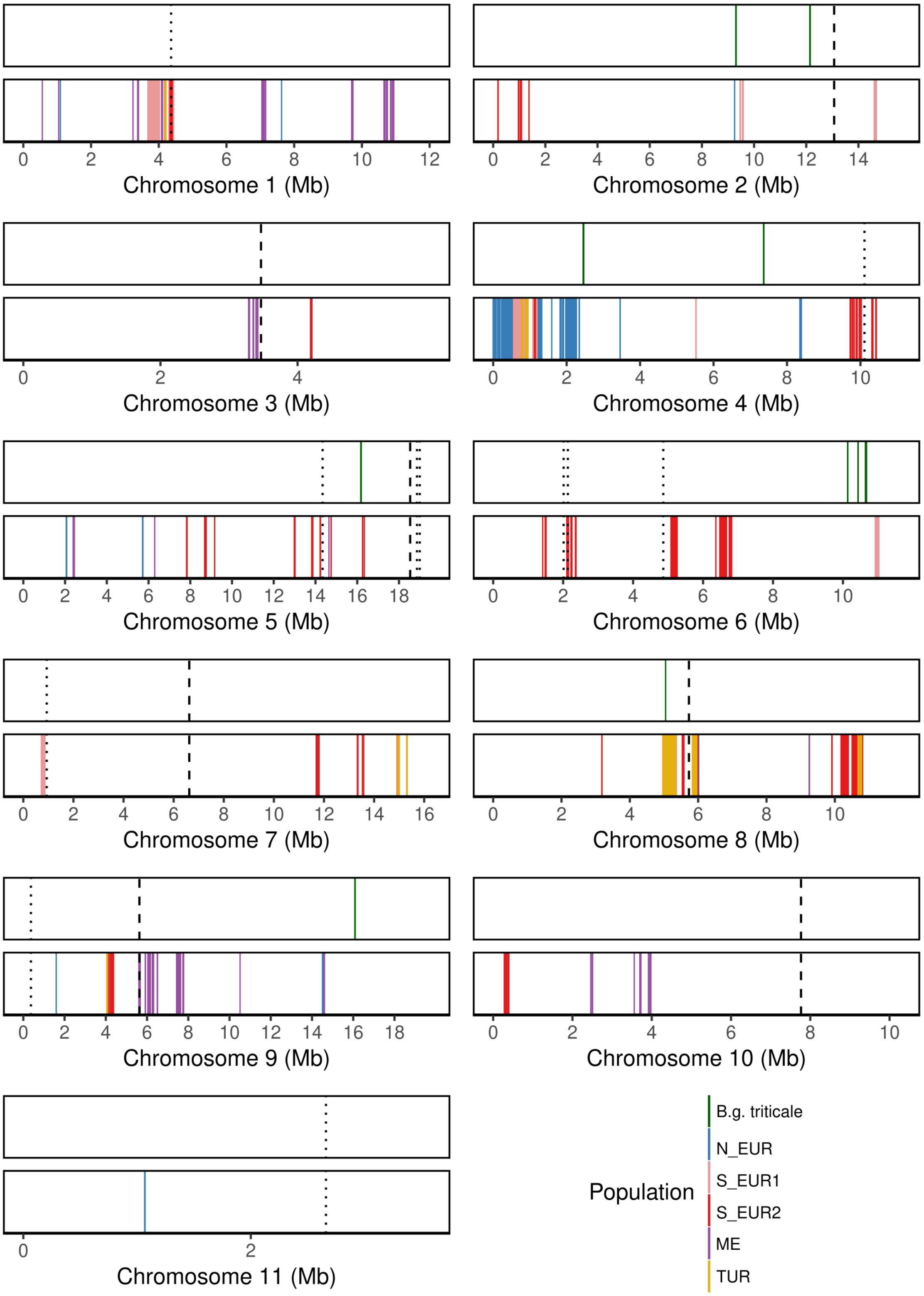


**Fig. S12. Comparing signatures of recent positive selection in *B.g. triticale* and *B.g. tritici*** Coloured vertical lines show the position of the SNPs associated with significant excess relatedness. Green lines in the top panels show SNPs crossing the False Discovery Rate (FDR) corrected 0.05 significance threshold for all *B.g. triticale* isolates. The bottom panels show regions under positive selection identified in Jigisha et al. 2025[1]). Such analysis was performed independently in each of the 5 *B.g. tritici* populations (ME, N_EUR, S_EUR1, S_EUR2, TUR. The dashed and dotted lines represent positions of known *B.g. tritici* avirulence genes and fungicide targets respectively (see Table S6).

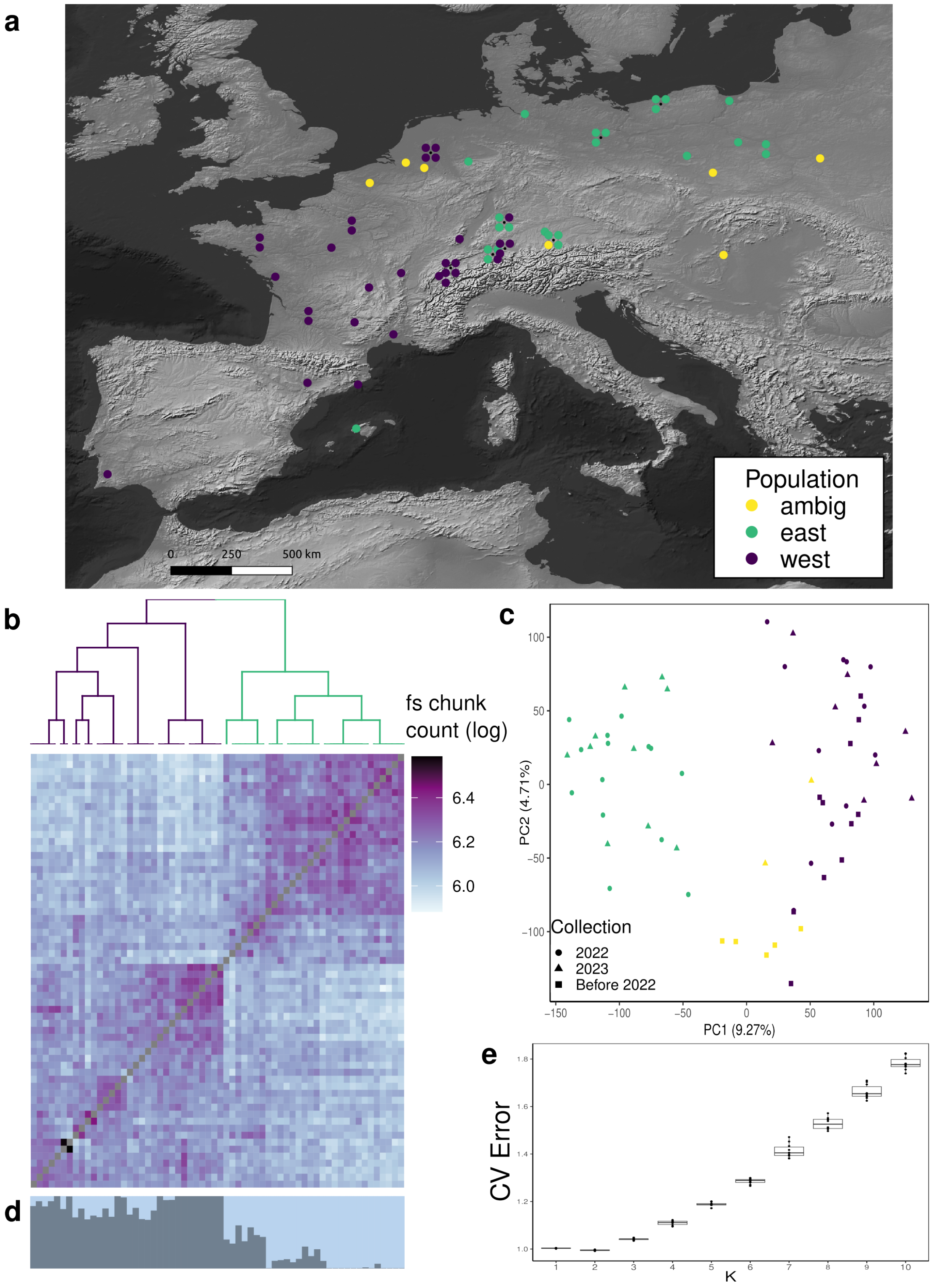


**Fig. S13. Population structure analysis of *B.g. triticale***
(**a**) Sampling locations of *B.g. triticale* isolates belonging to the east and west subpopulations. 'ambig' represents the 7 isolates that could not be assigned to either subpopulation conclusively (see S4 Appendix). (**b**) Dendrogram and coancestry matrix from the fineSTRUCTURE analysis of 62 *B.g. triticale* isolates. Higher chunk count reflects higher shared ancestry. (**c**) Principal component analysis of the 62 *B.g. triticale* isolates based on biallelic SNPs. Each point represents an isolate, coloured according to the population it was found to belong to (see S4 Appendix). (**d**) Ancestry proportions for all *B.g. triticale* isolates obtained from the ADMIXTURE analysis with K = 2. The order of isolates in panels b and d is the same. (**e**) Cross-validation errors of each of the 10 ADMIXTURE replicates performed for K from 1 to 10.

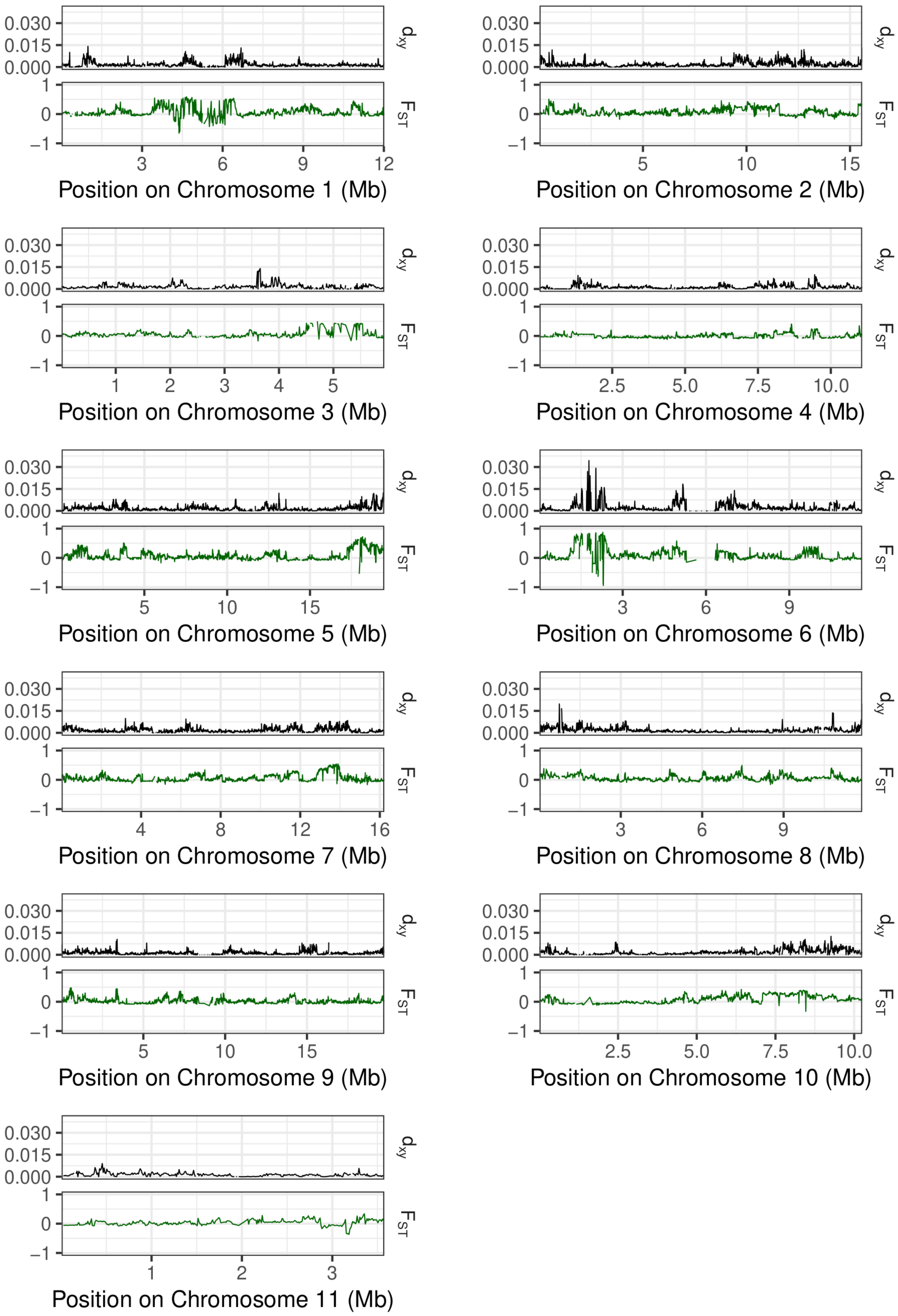


**Fig. S14. Genome-wide estimates of divergence between the eastern and western *B.g. triticale* populations**

$D_{xy}$ and $F_{ST}$ estimated between the eastern and western populations of triticale powdery mildew, along the genome in windows of 10 Kb.

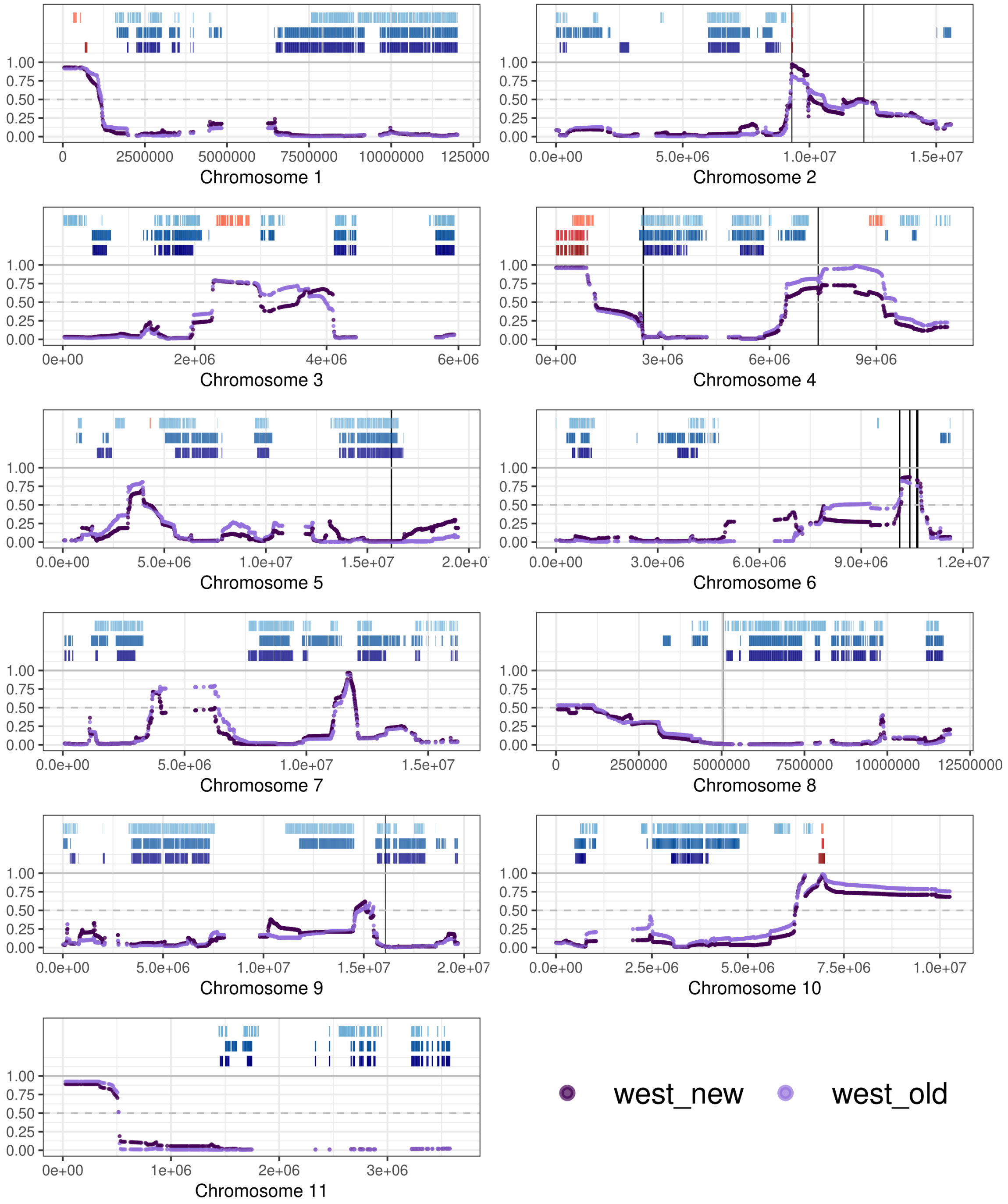


**Fig. S15. Variation in mean local ancestry over time**
The purple points show the mean *B.g. secalis* ancestry among the recently sampled (2022-2023; dark purple) and old (2009-2013; light purple) *B.g. triticale* isolates, calculated in windows of 10 Kb along the genome by MOSAIC using the new *B.g. tritici* donor panel. The three rows of segments in the upper-part of each panel show the regions fixed for *B.g. tritici* (blue) or *B.g. secalis* (red) ancestry in contemporary *B.g. triticale* populations. The first row from the top shows results from the substitution sites method, the second row corresponds to the MOSAIC run with old *B.g. tritici* donors and the third belongs to the MOSAIC run with new *B.g. tritici* donors. The vertical black lines represent loci found to be under recent positive selection from the isoRelate analysis, after FDR correction.

### Appendices

**Appendix S1. Looking for recent *B.g. secalis* introgression in *B.g. tritici***

If reproductive isolation between *B.g. triticale* and the parental *f. sp. B.g. tritici* was weak, we would expect to find signatures of recent introgression in *B.g. tritici.* Specifically, we would expect genomic segments originating from *B.g. secalis* in the *B.g tritici* population, transferred through sexual reproduction between *B.g. triticale and B.g. tritici.* The temporally staggered and spatially dense nature of sampling in our dataset from Europe allowed us to look for such introgressed regions in *B.g. tritici.*

We looked for *B.g. secalis* introgressions into the most recent *B.g. tritici* samples collected from 2022-2023. To do this we applied the substitution-sites method to look for *B.g. secalis* origin segments in *B.g. tritici.* We used 361,578 SNPs fixed for one allele in *B.g. secalis* and the other in *B.g. tritici* (substitution sites), and we considered individuals sampled until 2001, and therefore before the first detection of *B.g. triticale*. We assigned ancestry to genome segments of *B.g. tritici* individuals sampled from Europe in 2022 and 2023 (see Methods). These segments covered 70.3% of the whole genome, on average. *B.g. triticale*-mediated introgression would result in large segments of *B.g. secalis*-origin in the genomes of these *B.g. tritici* individuals. Overall, the *B.g. secalis* allele was present at 0.073-0.921% (mean = 0.284%) of the substitution sites across all 255 individuals, which corresponded to 0.004-0.36% (mean = 0.076%) of the genome-length covered by continuous segments. The proportion of the genome made up of *B.g. secalis*-origin segments differed between the five *B.g. tritici* populations (ANOVA, Df = 4, $p < 0.001$), with samples from the southern European population S_EUR2 showing the highest *B.g. secalis* contribution (Fig. S3a-b). We extracted all *B.g. secalis*-origin segments containing at least 10 substitution sites and found only 5 segments, three on chromosome 2 and two on chromosome 5, to be larger than 10 Kb in size (Fig. S3c). The three segments on chromosome 2 overlapped with each other and spanned ca. 14 Kb overall. These segments were found in 26 isolates overall, 24 of which belonged to the S_EUR2 population. The other two regions on chromosome 5 were only found in one isolate each (Fig. S3c). We consider these *B.g. secalis* segments to be a consequence of our panel of *B.g. tritici* sampled before the hybridization, which included only a small set of isolates from Northern Europe. Therefore these segments likely represent haplotypes that were shared between *B.g. tritici* and *B.g. secalis* before the emergence of *B.g. triticale*, but that were not captured in our small panel of isolates sampled before 2001. Nevertheless, the absence of widespread, large *B.g. secalis*-like segments in the recently sampled *B.g. tritici* individuals points towards a lack of recent, *B.g. triticale*-mediated introgression from *B.g. secalis* to *B.g. tritici.* The small number of *B.g. secalis* samples in our dataset prevented us from explicitly testing for introgression in the other direction, i.e. from *B.g. tritici* to *B.g. secalis.* However, given that rye powdery mildew and triticale powdery mildew rarely occur on the same host, it is unlikely that gene flow with *B.g. secalis* would be more frequent than with *B.g.*

*tritici*.

##### Appendix S2. Phylogenetic analysis of mating type genes

*Blumeria graminis* has two alternate mating types, 1-1 and 1-2. Successful sexual reproduction can take place only between two individuals with opposing mating types. The mating type locus of *B.g. tritici* (located on chromosome 1 in the reference genome 96224) contains either the *MAT1-1* or the *MAT1-2* genes, flanked on one side by *SLA2* and by *APN2/COX13* on the other [2, 3]. We performed a phylogenetic analysis for three of these genes (*MAT1-2* and *SLA2* associated with the 1-2 mating type, and *MAT1-1* associated with the 1-1 mating type) for 484 isolates of powdery mildew sampled from Europe and neighboring regions, including the three *formae speciales*- tritici, secalis and triticale.

In our dataset, 194 *B.g. tritici*, 35 *B.g. triticale* and 3 *B.g. secalis* isolates were of the 1-2 mating type (Fig. S5). There were three SNPs that differentiated *B.g. tritici* from *B.g. secalis* across the *MAT1-2* and *SLA2* genes. All *B.g. triticale* isolates with this mating type showed the *B.g. tritici* genotype at these positions. The remaining 250 individuals, including 4 *B.g. secalis*, 27 *B.g. triticale* and 219 *B.g. tritici*, were of the 1-1 mating type. The *MAT1-1* gene had 12 SNPs differentiating *B.g. secalis* from *B.g. tritici*. Both the *B.g. secalis* and the *B.g. tritici* haplotypes could be found among the *B.g. triticale* samples. Menardo and colleagues [4], using a subset of the isolates used in this study, also found similar results where *B.g. triticale* individuals exclusively showed the *B.g. tritici* haplotype for one mating type. The authors concluded that one idiomorph of the mating type gene was inherited from *B.g. secalis* during the initial hybridization event, and the other idiomorph was brought into the *B.g. triticale* gene pool through subsequent backcrossing with *B.g. tritici*. Ongoing gene flow between *B.g. secalis* and *B.g. triticale* could potentially result in the introgression of the MAT1-1 idiomorph from *B.g. secalis* into the recent hybrid. Since we did not detect this, we cannot confirm that there have been new instances of hybridization or backcrosses with *B.g. secalis* in the recent past.

##### Appendix S3. Inferring local ancestry in *B.g. triticale* genomes

In addition to assigning parental origin to genome segments using substitution sites, we also performed local ancestry inference in *B.g. triticale* using a probabilistic approach, as implemented in MOSAIC [5]. This method infers the relationship between ancestral admixing groups and candidate donor panels from the data and therefore allows for the donors to be descendants of the unseen admixing groups. We modelled *B.g. triticale* genomes to be the result of a 2-way admixture event using two different sets of donor panels, ran as two independent iterations of MOSAIC. In the first, we included 7 *B.g. secalis* isolates and 17 *B.g. tritici* isolates that had been sampled from Europe until 2001. This donor panel was the same as the one used in the substitution-sites analysis. In the second run of MOSAIC, we used as donors the same set of *B.g. secalis* individuals as above, but replaced the old *B.g. tritici* panel with recently sampled wheat powdery mildew (collected in 2022-2023) that belonged to the Northern European (N_EUR) population. For both runs, MOSAIC inferred the two ancestries to closely resemble *B.g. tritici* and *B.g. secalis* (Table S4).

MOSAIC computes local ancestry along the genome at grid-points evenly spaced on genetic distance. These can be converted to ancestry estimates at physical SNP positions. We then divided the genome (chromosomes 1-11) into non-overlapping 10 Kb windows, and obtained 9919 windows with at least 1 SNP. For each isolate, we then calculated the average ancestry of all SNPs within each of the windows. If the mean probability of *B.g. secalis* ancestry of a window was greater than 0.7, the window was assigned to be of *B.g. secalis* ancestry in that isolate, and if it was less than 0.3, then the window was assigned to *B.g. tritici* ancestry. We also used the substitution-sites approach of assigning SNPs to parental origin to obtain mean ancestry in the 10 Kb windows (Methods). Here, the ancestry associated with each SNP was either *B.g. tritici B.g. secalis* based on their allele. Under this approach, we considered a window to be of *B.g. secalis* ancestry in an isolate if all SNPs in that window could be assigned to *B.g. secalis* and viceversa the window was determined to be of *B.g. tritici* ancestry if all SNPs were assigned to *B.g. tritici.*

To evaluate genome stability, we identified regions that were fixed for an ancestry in *B.g. triticale.* These were the windows that were assigned the same ancestry across all 46 *B.g. triticale* isolates sampled in 2022-2023. Between a third to a half of the genome was found to be fixed for one ancestry or another, depending on the method used (Table S5). Regions fixed for *B.g. secalis* made up a small proportion of this, and were restricted to a few chromosomes (Fig. 3, Figs. S8-S9). The results across both methods of MOSAIC were largely consistent. The substitution–sites approach detected more *B.g. secalis-fixed* regions than MOSAIC, but these regions were generally inferred to have high mean *B.g. secalis* ancestry by MOSAIC as well (Fig. 3, Figs. S8-S9). Overall, we found that a substantial proportion of the genome of this hybrid has nearly stabilized within 25 years of its emergence.

##### Appendix S4. Population structure analyses of *B.g. triticale*

The fineSTRUCTURE analysis of all European powdery mildew isolates in our dataset indicated that *B.g. triticale* populations in Europe were structured. To explore this further, we performed a principal component analysis (Fig. S13) using 536,386 biallelic SNPs across all 62 non-clonal triticale mildew isolates. PC1 explained 9.27% of the total variation and roughly separated individuals sampled from the eastern part of the sampling range from those sampled from the west. The ADMIXTURE analysis also revealed the best K to be 2 and suggested a similar east-west division (Fig. S13). We repeated the fineSTRUCTURE analysis with just the *B.g. triticale* isolates (Fig. S13), and compared the results to the previous run with the three *formae speciales.* The east-west population classification was consistent across both fineSTRUCTURE runs for 55 of 62 triticale mildew isolates: 31 individuals belonged to the western population and 24 to the eastern population. The 7 remaining isolates were inconsistently grouped in different populations with different methods and were therefore excluded from between-population comparisons (Fig. S13). The two populations showed low levels of differentiation generally, with median $F_{ST} = 0.02525$ and median $D_{xy} = 0.00125$.
